# Imbalance between IL-36 receptor agonist and antagonist drives neutrophilic inflammation in COPD

**DOI:** 10.1101/2021.10.06.463311

**Authors:** Jonathan R Baker, Peter S Fenwick, Carolin K Koss, Harriet B Owles, Sarah L Elkin, Matthew Thomas, Jay Fine, Karim C El Kasmi, Peter J Barnes, Louise E Donnelly

## Abstract

Current treatments fail to modify the underlying pathophysiology and disease progression of chronic obstructive pulmonary disease (COPD), necessitating novel therapies. Here, we show that COPD patients have increased IL-36γ and decreased IL-36 receptor antagonist (IL-36Ra) in bronchoalveolar and nasal fluid compared to control subjects. IL-36γ is derived mainly from small airway epithelial cells (SAEC) and further induced by a viral mimetic, whereas IL-36RA is derived from macrophages. IL-36γ stimulates release of the neutrophil chemoattractants CXCL1 and CXCL8, as well as elastolytic matrix metalloproteinases (MMPs) from small airway fibroblasts (SAF). Proteases released from COPD neutrophils cleave and activate IL-36γ thereby perpetuating IL-36 inflammation. Transfer of culture media from SAEC to SAF stimulated release of CXCL1, that was inhibited by exogenous IL-36RA. The use of a therapeutic antibody that inhibits binding to the IL-36 receptor (IL-36R) attenuated IL-36γ driven inflammation and cellular cross talk. We have demonstrated a novel mechanism for the amplification and propagation of neutrophilic inflammation in COPD and that blocking this cytokine family via a IL-36R neutralising antibody could be a promising new therapeutic strategy in the treatment of COPD.

## Introduction

Chronic obstructive pulmonary disease (COPD) is a major global health burden which affects ∼10% of people over 45 years of age and is the 3rd commonest cause of death in the world (Collaborators, 2020). COPD is a chronic inflammatory lung disease, associated with increased numbers of inflammatory cells, including macrophages, neutrophils and lymphocytes (Barnes, 2016b). Inflammation occurs predominantly in the lung parenchyma and peripheral airways and results in irreversible and progressive airflow limitation due to small airway fibrosis and parenchymal destruction (emphysema). Current therapies targeting the inflammatory mediators of COPD have been poorly effective clinically and do not reduce disease progression, so novel anti-inflammatory approaches are greatly needed (Barnes, 2019).

Interleukin (IL)-36 cytokines belong to the IL-1 superfamily and comprise three receptor agonists (IL-36α, IL-36β and IL-36γ) and two receptor antagonists (IL-36Ra and IL-38). The IL-36 receptor comprises a heterodimer of IL-36R and the IL-1 receptor accessory protein (IL-1RAcP), so that IL-36R competes with the IL-1R for IL-1RAcP (Gresnigt and van de Veerdonk, 2013). Binding of IL-36 agonists to the receptor leads to the recruitment of IL-1RAcP and subsequent down-stream signalling via nuclear factor (NF)-КB and mitogen-activated protein kinase (MAPK) pathways (Bassoy et al., 2018). IL-36 cytokines may therefore play an important role in the chronic release of pro-inflammatory cytokines and chemokines in inflammatory conditions, such as COPD.

IL-36α, IL-36β, IL-36γ and IL-36Ra are all secreted in an inactive form and require N-terminal cleavage by serine proteases to induce a 500-fold increase in their activity (Henry et al., 2016). IL-36 isoforms are cleaved by neutrophil products, specifically neutrophil elastase, proteinase-3 and cathepsin G, with each exhibiting selectivity (Clancy et al., 2018). Neutrophils are markedly increased in the lungs and airways of patients with COPD and are associated with increased secretion of several proteases (Stockley, 1999); which are further increased during acute exacerbations (Barnes, 2016a; Gompertz et al., 2001). In COPD, this could lead to persistent activation of IL-36 and subsequent downstream inflammatory signalling.

Elevated levels of IL-36 cytokines been reported in COPD, with IL-36α and IL-36γ being elevated in plasma and bronchoalveolar fluid (BALF) from smokers with and without COPD compared to healthy controls, although in a small population of only 5 patients with mild disease (Kovach et al., 2020). IL-36γ is also increased in sputum samples from COPD patients with neutrophilic phenotype (Li et al., 2021). A similar association of IL-36γ with neutrophilia has been reported in patients with obstructive lung disease, where as a decrease in IL-36γ associated with eosinophilia (Moermans et al., 2021). As well as the importance of examining the levels of receptor agonists, a recent study suggests there is a reduction in IL-36Ra in both the serum and the sputum of pediatric asthmatic patients (Liu et al., 2020). These authors also showed that intra-nasal administration of IL-36Ra in a murine model of asthma reduced airway hyperresponsiveness and inflammatory cell infiltrates into the lung (Liu et al., 2020).

Most of the current understanding of the role of IL-36 in respiratory disease has come from *in vivo* mouse models of lung disease. Intra-tracheal instillation of *Pseudomonas aeruginosa* in mice increased expression of IL-36α and IL-36γ in bronchoalveolar lavage (BAL) fluid and lung homogenates (Aoyagi et al., 2017a). Subsequent *in-vitro* experiments confirmed that this was due to increased expression by alveolar macrophages and epithelial cells (Aoyagi et al., 2017b). Of note, knockout of *either IL-36R* or *IL-36γ*, but not *IL-36α*, protected against lung damage and increased survival time caused by infection of this bacterium (Aoyagi et al., 2017b). By contrast, when mice were exposed to an intranasal influenza viral challenge, there was induction of IL-36α, but not IL-36γ protein in the lung, although *IL-36γ* mRNA was induced (Aoyagi et al., 2017a). Others have suggested that IL-36γ is up-regulated following viral infection and is a protective mechanism due to skewing of macrophage phenotypes towards a more pro-resolving M2 phenotype (Wein et al., 2018). We have recently shown that intratracheal delivery of IL-36γ into the lung of mice increases neutrophil chemokines and numbers. Using a COPD exacerbation model of cigarette smoke exposure in combination with influenza H1N1, we showed that IL-36R deficient mice have reduced neutrophil recruitment into their lungs and inflammatory mediators (Koss et al., 2021).

Studies using human airway cells have shown that bronchial epithelial cells stimulated with the viral mimetic, dsRNA, show increased expression of IL-36γ that was further enhanced by IL-17 (Chustz et al., 2011). IL-36γ, but not IL-36α, was also induced in human peripheral blood mononuclear cells (PBMC) by the fungus *Aspergillus fumigatus* (Gresnigt et al., 2013). This is important as 37% of stable COPD patients are colonised with *A. fumigatus* (Bafadhel et al., 2014), potentially due to an inability of COPD macrophages to phagocytose these fungal spores (Wrench et al., 2018). IL-36α, β and γ all induce protein and gene expression of IL-6 and CXCL8 in normal human lung fibroblasts and bronchial epithelial cells, whereas exogenous addition of IL-36Ra reduces these mediators (Zhang et al., 2017). This suggests that IL-36 cytokines may induce the release of these proinflammatory cytokines via NF-κB and MAPK signalling (Zhang et al., 2017).

Elevated concentrations of IL-36 cytokines have been associated with a variety of chronic inflammatory diseases, including generalized pustular psoriasis (GPP), inflammatory bowel disease, rheumatoid arthritis, and systemic lupus erythematosus. Loss of function mutations within the IL-36Ra gene (*IL-36RN*) may result in GPP, an inflammatory skin disease characterized by elevated proinflammatory cytokines and immune cell and neutrophil infiltrates (Marrakchi et al., 2011). Furthermore, there are promising data emerging from a completed phase 1 clinical trial using a neutralising antibody targeting IL-36R in this disease (Bachelez et al., 2019). However, the role of the IL-36 family of cytokines has not been explored in COPD.

In the current study, we hypothesized that there is an altered expression of IL-36 cytokines in the COPD lung and this imbalance drives the characteristic neutrophilic inflammation seen in this disease. We therefore examined the expression of IL-36 agonist and antagonist cytokines in the lungs of COPD patients compared to healthy individuals. We studied the cellular source of IL-36 within the lung and identified the effector cells of this family of proinflammatory cytokines. We also examined the mechanism by which IL-36 may amplify neutrophilic inflammation in the lungs of COPD patients and highlight a critical role for these cytokines in perpetuating neutrophilic inflammation and progression in COPD.

## Results

### Increased IL-36 cytokines in COPD

Firstly, IL-36 cytokines were examined in BAL fluid from age-matched non-smokers, smokers and COPD patients. Elevated concentrations of IL-36γ were found in both smoker and COPD patients compared to non-smokers (Fig. 1A), whereas IL-36α and IL-36β were not detected (Supplementary Fig. 1A, B). Increased release of IL-36γ was also measured in nasal secretions from COPD patients compared with control subjects (Fig. 1B) with concentrations being much higher (ng/ml) than the diluted BAL fluid (pg/ml). No changes in IL-36α or IL-36β were observed in nasal fluid samples (Supplementary Fig. 1C, D), suggesting that IL-36γ is specifically up-regulated in the airway mucosa. IL-36 cytokines are also reported to be increased in the serum (Kovach et al., 2020) and sputum of COPD patients (Li et al., 2021). However, we found no significant increase in IL-36γ, in sputum samples from smokers or COPD patients (Fig. 1C), although it was detectable and trended towards an increase. Neither IL-36α nor IL-36γ were altered in COPD serum samples, with most below the level of detection of the assay (Supplementary Figure 1E and F). Gene expression was also examined in lung homogenate samples from non-smokers, smokers and COPD patients. *IL-36γ* mRNA was significantly up-regulated in COPD patients (Fig. 1D).

**Figure 1.**
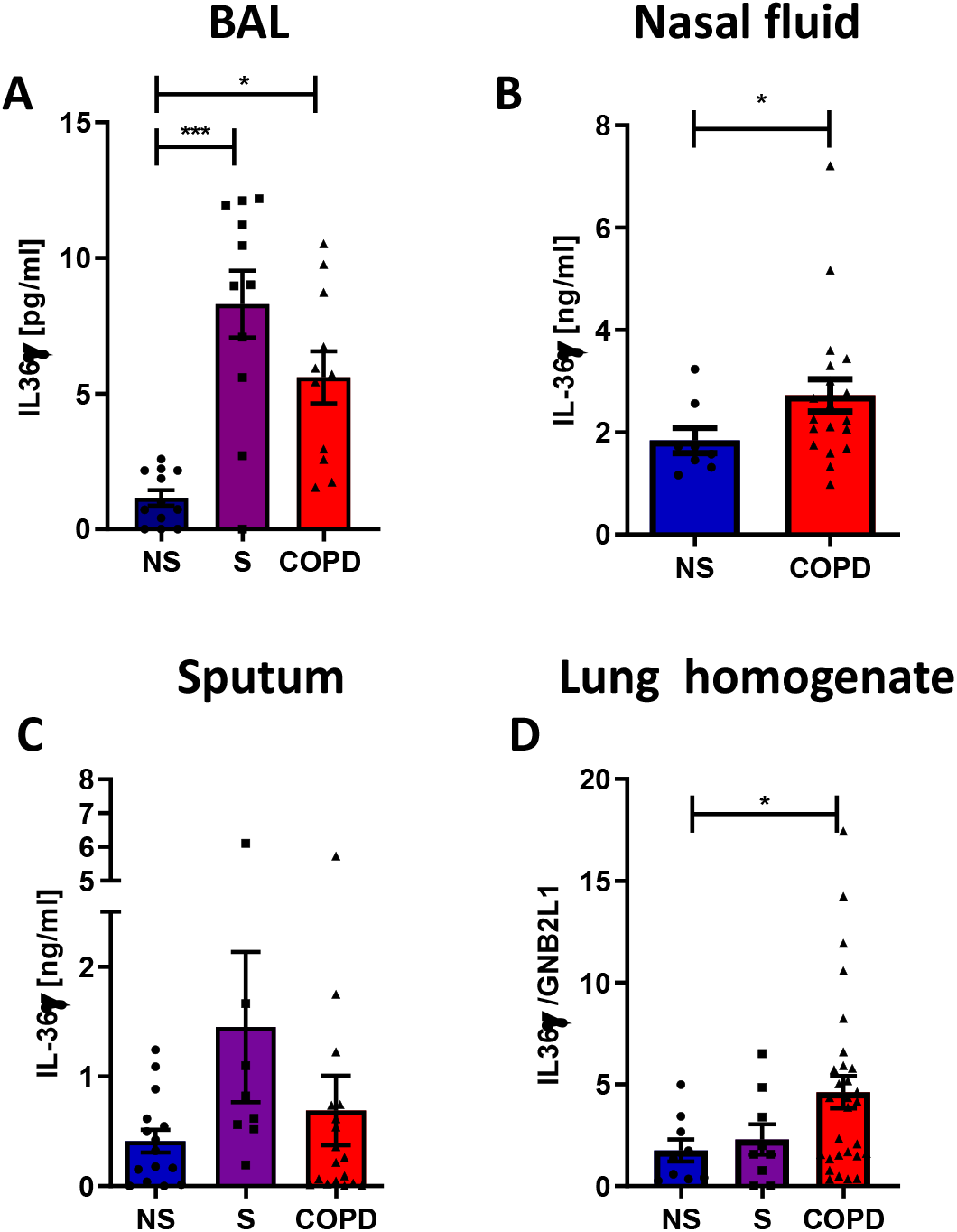
IL-36γ protein is elevated in COPD. A) IL-36γ was measured in the bronchoalveolar lavage fluid (BAL) of non-smokers (NS, ● Blue, n=12), smokers (S, ■ purple, n=11) and COPD patients (▲, red, n=11); B) nasal fluid of NS (n=8) and COPD patients (n=20) and C) sputum of NS (n=15), smokers (n=8) and COPD patients (n=18) by ELISA. IL-36γ gene expression was examined in D) lung homogenate samples from NS (n=9), smokers (n=9) and COPD patients (n=29). Data are means ± SEM and analysed by Kruskal-Wallis test with post-hoc Dunn’s test or by Mann-Whitney U test; * P <0.05, ***P<0.001.

### Cellular source of IL-36 in the lung

To determine the cellular source of elevated IL-36γ within the airways of COPD patients, gene expression of IL-36 cytokines was examined in key cells involved in COPD pathogenesis: lung tissue-derived macrophages (TMφ), small airway fibroblasts (SAF) and small airway epithelial cells (SAEC). TMφ from COPD patients displayed a non-significant trend towards increased *IL-36γ* expression in cells from smokers and COPD patients compared with controls (Fig. 2A). Expression of *IL-36α* was unchanged between patient groups, whilst *IL-36β* RNA was significantly down-regulated in smokers but unchanged in COPD patients compared to non-smokers (Supplementary Fig. 2A and 2B). *IL-36α* expression was not detected in SAF and there was no difference in gene expression of *IL-36γ* between COPD SAF and non-smokers (Fig. 2B), with a similar pattern for *IL-36β* (Supplementary Fig. 2C). In contrast, *IL-36γ* expression was significantly increased in COPD SAEC compared to non-smokers (Fig. 2C), with a significant increase in *IL-36α* expression but no change in *IL-36β* expression (Supplementary Fig. 2D and E). IL-36γ protein release from TMφ and SAF was below the level of detection. However, IL-36γ release from SAEC was detectable and COPD SAEC released significantly more IL-36γ compared to non-smoking controls (Fig. 2D), suggesting that airway epithelial cells could be a major source of IL-36γ in peripheral airways, under basal conditions, and that these cells may be responsible for the elevated levels observed in BAL fluid and nasal secretions in COPD.

**Figure 2.**
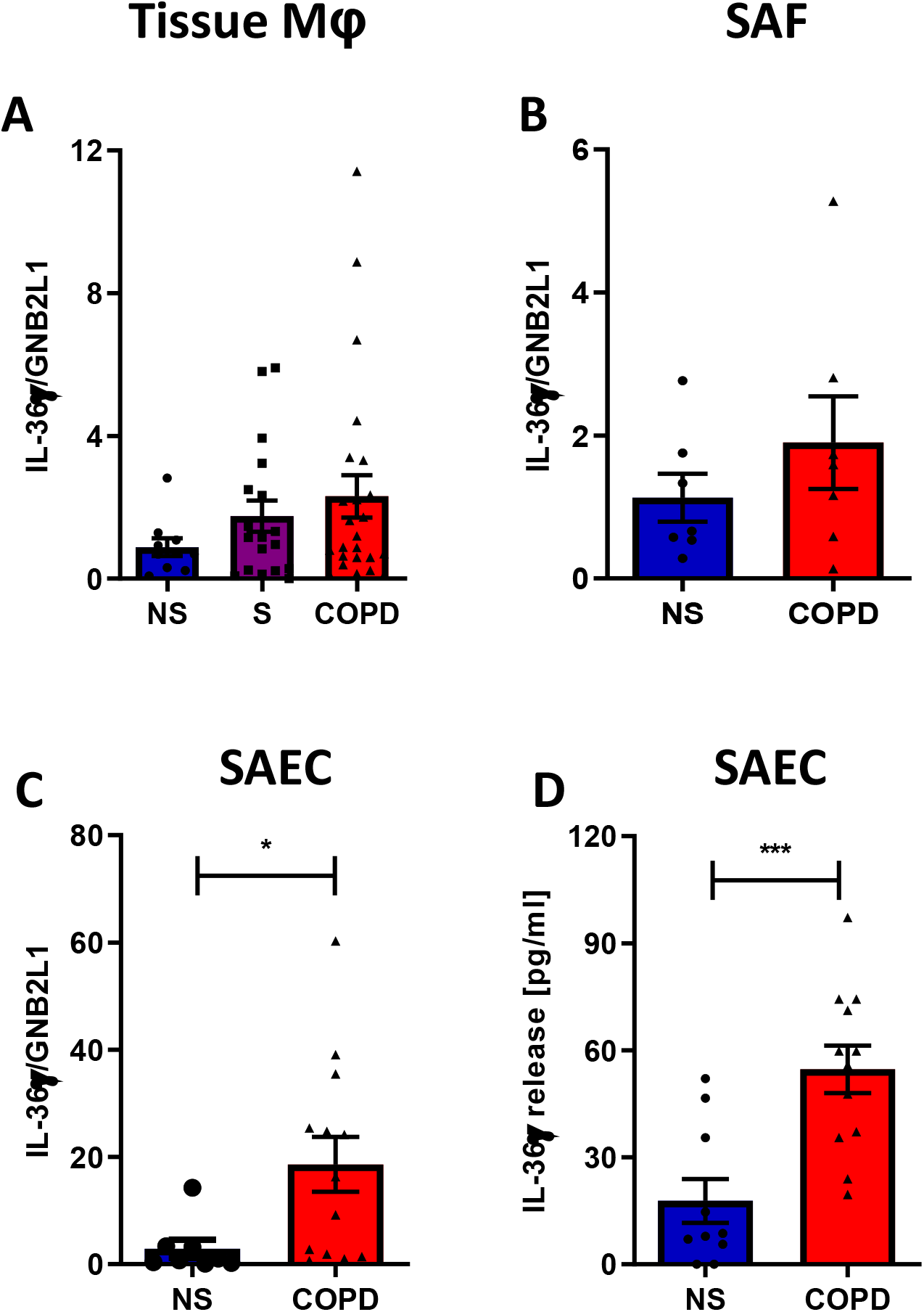
Small airway epithelial cells are the source of IL-36γ in COPD. Gene expression of *IL-36γ* was measured in A) lung tissue-derived macrophages (TMφ), B) small airway fibroblasts and C) small airway epithelial cells from non-smokers (NS, ● n=5-10), smokers (S, ■ n=7-18) and COPD patients (▲, n=7-24). IL-36γ release from H) small airway epithelial cells from NS (n=10) and COPD (n=12) patients, measured by ELISA. Data are means ± SEM and analysed by Kruskal-Wallis test with post-hoc Dunn’s test or by Mann-Whitney U test; * P <0.05, ***P<0.001.

### IL-36Ra is reduced in COPD

IL-36 signalling can be modulated by binding of IL-36Ra to the IL-36R, thereby supressing receptor activation (Bassoy et al., 2018). Genetic mutations in the IL-36Ra can drive disease, with a loss of function mutation in IL-36Ra being a likely mechanism in GPP (Marrakchi et al., 2011). These data suggest not only the importance of the up-regulation of IL-36 agonists, but also modulation of the IL-36 receptor antagonist. The levels of IL-36Ra in the context of COPD have not previously been reported.

The protein levels of IL-36Ra were significantly reduced in BAL and sputum of COPD patients compared to non-smokers (Fig. 3A and B). IL-36Ra gene expression (*IL-36RN*) was also significantly reduced in COPD lung homogenate samples compared to non-smokers and smokers (Fig. 3C), with smokers showing a trend towards increased *IL-36RN* expression, potentially as a protective mechanism.

**Figure 3.**
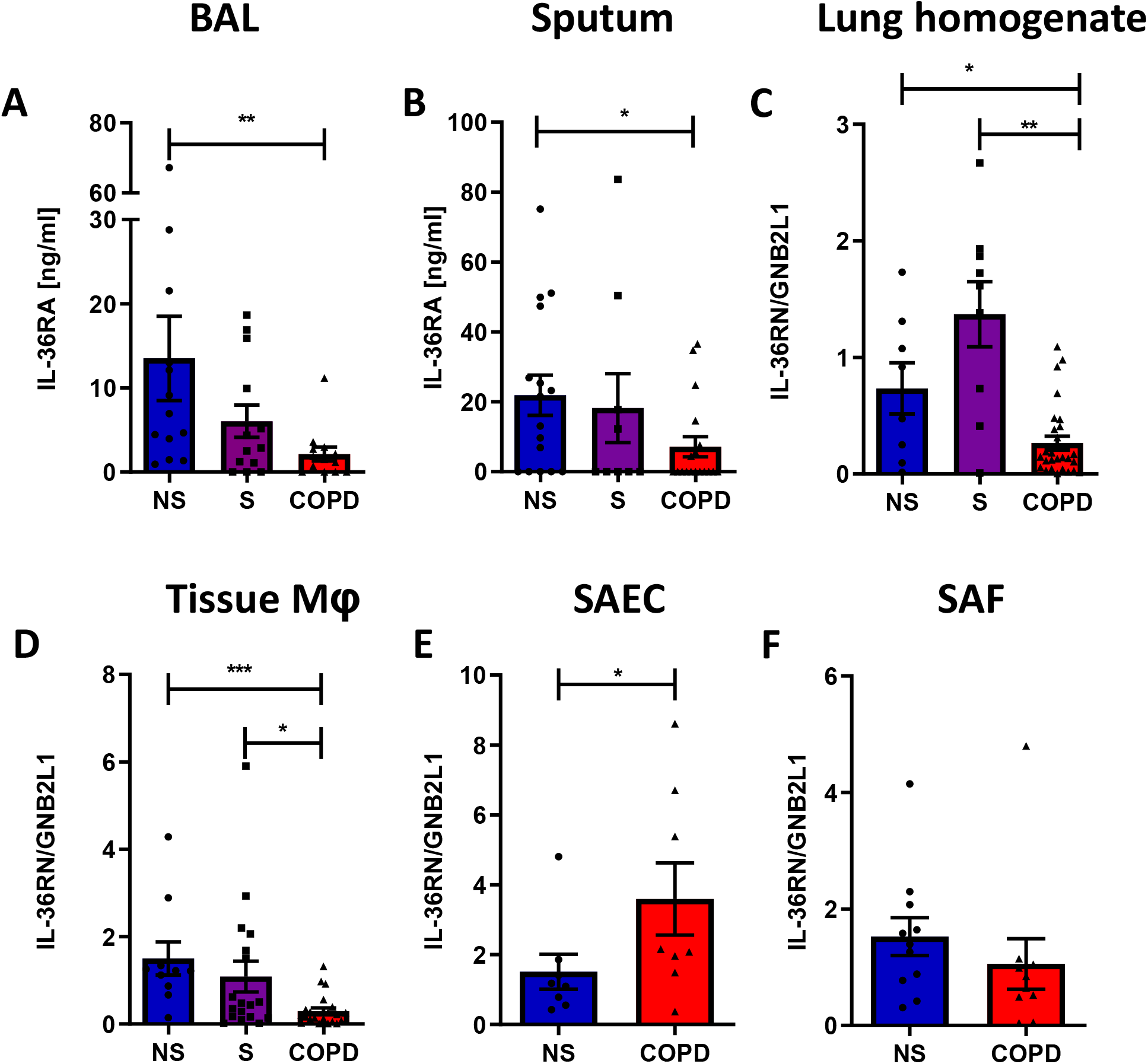
IL-36 receptor antagonist is reduced in COPD patients. IL-36Ra protein was detected in the A) broncholalveolar lavage fluid and B) sputum of non-smokers (NS, ● n=13-16), smokers (S, ■ n=9-13) and COPD (▲ n=13-18) patients by ELISA. Gene expression of IL-36RN was detected in C) Lung homogenate samples from NS (n=9), smokers (n=9), COPD patients (n=29), D) lung tissue-derived macrophages, E) small airway epithelial cells and F) small airway fibroblasts S (n=5-10), smokers (n=7-18) and COPD patients (n=7-24). Data are means ± SEM and analysed by Kruskal-Wallis test with post-hoc Dunn’s test or by Mann-Whitney U test; * P <0.05 ** P <0.01.

To determine the cellular source of IL-36Ra, we examined gene expression of *IL-36RN* in different cells. *IL-36RN* was significantly down regulated in TMφ from COPD patients compared to both non-smoker and smoker samples (Fig. 3D), this appeared to be specific for the receptor antagonist of IL-36 as this was not seen for IL-1 receptor antagonist (Supplementary Fig. 2F). SAEC displayed a significant increase in *IL-36RN* in cells from COPD patients (Fig. 3E), whilst there was no change in expression of *IL-36RN* in SAF between non-smoker and COPD patients (Fig. 3F). Gene expression of the other IL-36 antagonist, IL-38, was undetectable in all cell types examined. As changes in IL-36R expression may also alter IL-36 signalling, the gene expression of the IL-36 receptor was examined and found to be unchanged in all patient groups, in TMφ, SAEC and SAF (Supplementary Fig. 2G-H). These data suggest that along with elevated levels of IL-36γ in COPD patients, there is dysregulation of the receptor antagonist which may amplify the effects of IL-36 on the lung.

### IL-36γ induction

As SAEC appear to be the main source of IL-36γ, we investigated whether SAEC could be stimulated to release more IL-36γ in response to stimuli which may exacerbate COPD. We showed that many of the common airway epithelium stimulants, such as cigarette smoke extract (CSE), TNFα, or the common colonising respiratory pathogen, *Haemophilus influenzae* failed to stimulate IL-36γ release (Fig. 4A). However, the TLR3 agonist, poly I:C stimulated release of IL-36γ from SAEC (Fig. 4A) and in a concentration-dependent manner (Supplementary Fig. 3A) with cells from COPD patients releasing more because of a higher baseline release of this cytokine. However, poly I:C failed to induce further release of IL-36α (Supplementary Fig. 3B) and IL-36β protein was undetectable in these cells. The same stimuli were applied to non-smoker and COPD SAF, with non-smoker SAF being unresponsive to all stimuli, whilst COPD SAF released more IL-36γ in response to all stimuli, although these were at a lower level when compared to SAEC (Supplementary Fig. 3C). These data suggest that COPD SAEC may respond to viral infection by releasing IL-36γ, thereby enhancing perpetuating the already elevated levels of IL-36γ in the lung.

**Figure 4.**
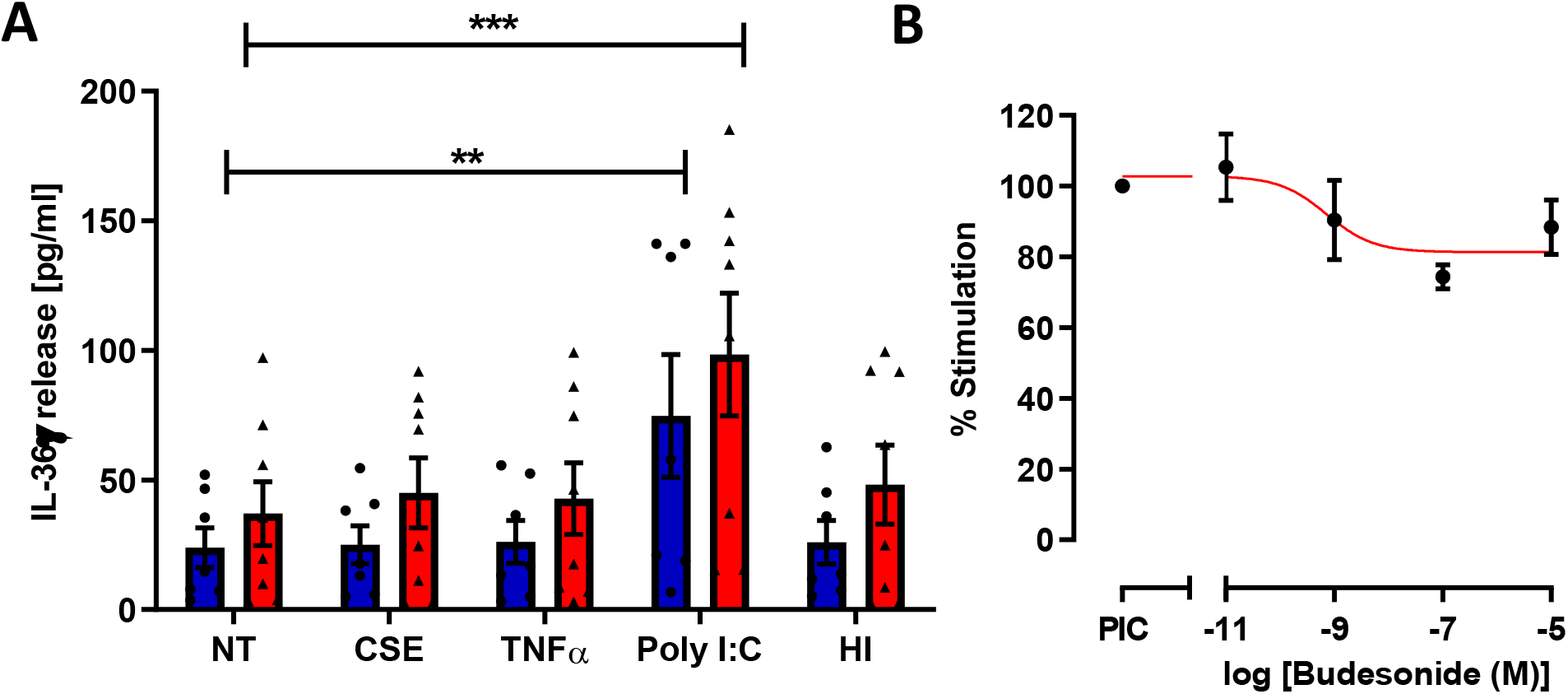
IL-36γ protein release is driven by steroid insensitive TLR3 activation. A) Small airway epithelial cells from non-smokers (blue bars, ● n=7) and COPD (red bars, ▲, n=8) patients were exposed to media alone (no treatment: NT), 10% (v/v) cigarette smoke extract (CSE), 10 ng/ml TNFα, 100μg/ml poly I:C, or 1.5×10^10^ CFU/ml *H. influenzae* (HI) for 24h. Media was collected and IL-36γ release measured by ELISA. B) Small airway epithelial cells from n=4 non-smokers were treated with 100 μg/ml poly I:C in the presence or absence of increasing concentrations of budesonide for 24h. Media was collected and IL-36γ was measured by ELISA. Data are means ± SEM and analysed by two way anova with post-hoc Dunnett’s multiple comparisons test; ** P <0.01, ***P<0.001 *vs*. NS. ^##^ P <0.01, ^###^ P<0.001 *vs*. COPD.

### Effect of glucocorticosteroids

As glucocorticosteroids are often given as an anti-inflammatory treatment to many COPD patients, particularly during viral-induced exacerbations, we examined whether TLR3 induction of IL-36γ was glucocorticosteroid-sensitive. However, increasing concentrations of the glucocorticosteroid, budesonide, had little effect on poly I:C-stimulated release of IL-36γ from SAEC (Fig.4B). These data suggest that treatment of a viral exacerbation with corticosteroids will not supress the elevated levels of IL-36γ that may be induced during a viral infection.

### Effects of IL-36γ on lung cells

IL-36γ activates multiple different cell types (Bassoy et al., 2018). Therefore, we investigated which pulmonary cells are the likely targets in COPD. TMφ, SAEC and SAF from COPD patients and controls were stimulated with activated IL-36γ and release of CXCL1 (GRO-α), CXCL8 (IL-8), IL-6 and granulocyte-macrophage colony-stimulating factor (GM-CSF) were measured, as they are all are increased in COPD lungs. Non-smoker, smoker and COPD TMφ were unresponsive to IL-36γ, with no change in CXCL8 or IL-6 release (Fig. 5A, B). Stimulation of SAEC with IL-36γ, led to a modest increase in both CXCL8 and IL-6 (Fig. 5C and D). However, there was a significant increase in CXCL1 release from both non-smoker and COPD SAEC (Fig. 5E).

**Figure 5.**
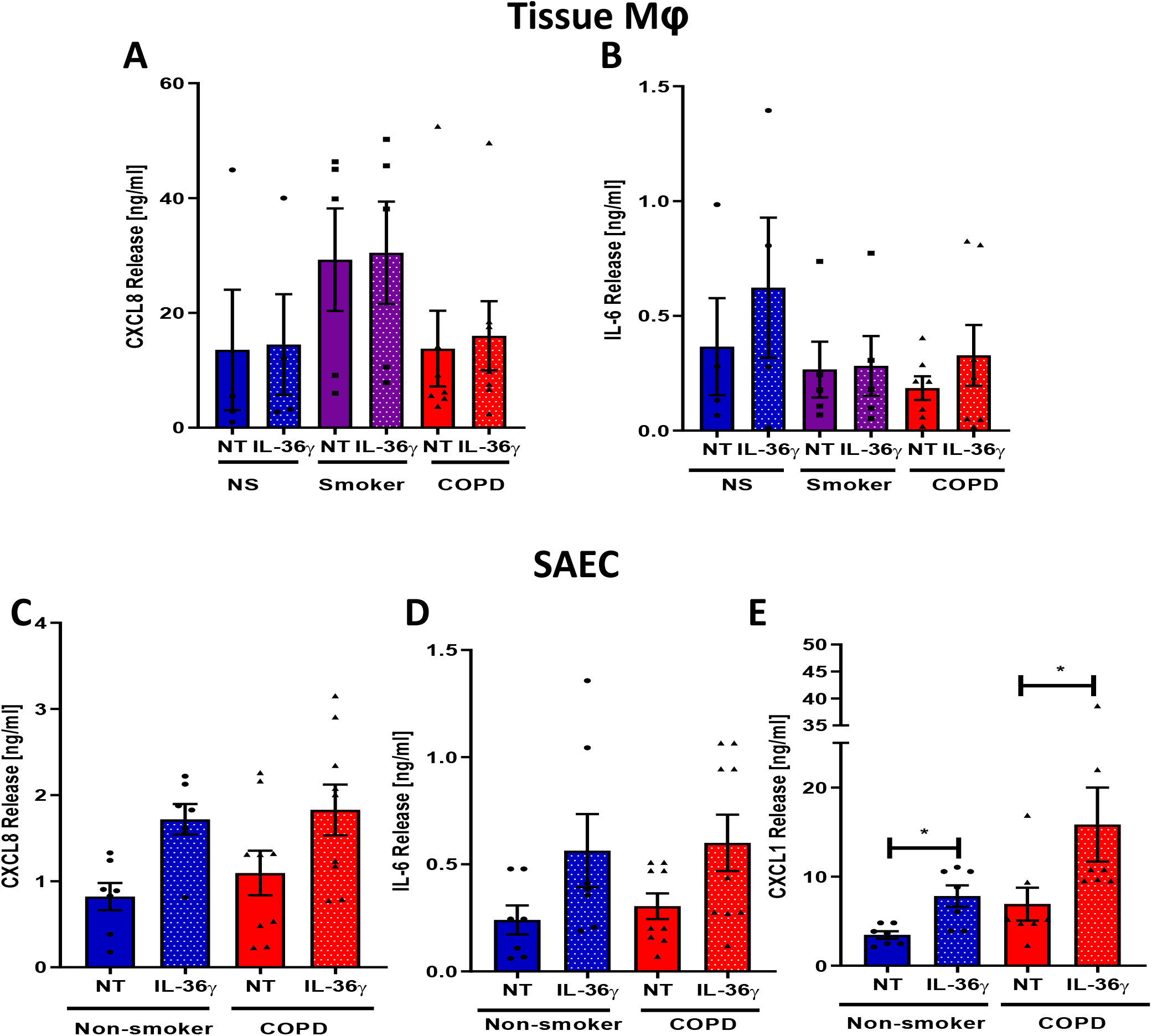
Effect of IL-36γ on lung tissue macrophages and small airway epithelial cells. Lung tissue-derived macrophages from non-smokers (n=4), smokers (n=5) or COPD (n=7) patients were incubated in the absence (NT) or presence of 100 ng/ml IL-36γ for 24h. Media was harvested and release of A) CXCL8 and B) IL-6 were measured by ELISA. Small airway epithelial cells from non-smokers (n=7) or COPD (n=7-9) patients were incubated in the absence (NT) or presence of 100 ng/ml IL-36γ for 24h. Media was harvested and release of C) CXCL8, D) IL-6 and E) CXCL1 were measured by ELISA. Data are means ± SEM and analysed by Kruskal-Wallis test with post-hoc Dunn’s test; * P <0.05, **P<0.01.

In contrast to the data derived from TMφ and SAEC, stimulation of SAF with IL-36γ induced a large and significant increase in the release of CXCL8, IL-6, CXCL1 and GM-CSF (Fig. 6A-D), with no difference between control and COPD cells. The magnitude of cytokine release was some 25-fold higher than from SAEC. These data strongly suggest that SAF are a key effector cell type for IL-36γ in the airways. Furthermore, these proinflammatory cytokines and chemokines are elevated in COPD patients (Barnes, 2009; Schulz et al., 2003; Traves et al., 2002) and may be responsible for neutrophil recruitment into the airways, which is a key feature of this disease. To examine whether these effects were specific to IL-36γ or if there was an additive effect of all three cytokines in combination, all cell types were treated with the individual IL-36 isoforms or in combination. CXCL8 release from SAF, SAEC and TMφ, was the same for each isoform and when in combination, with SAF still releasing the greatest levels of cytokines in response (Supplementary Fig 4A-C). These data again strongly suggest that SAF are the effector cell type for IL-36γ and may have a key role in neutrophil recruitment observed in this disease via increased neutrophil chemokine release.

**Figure 6.**
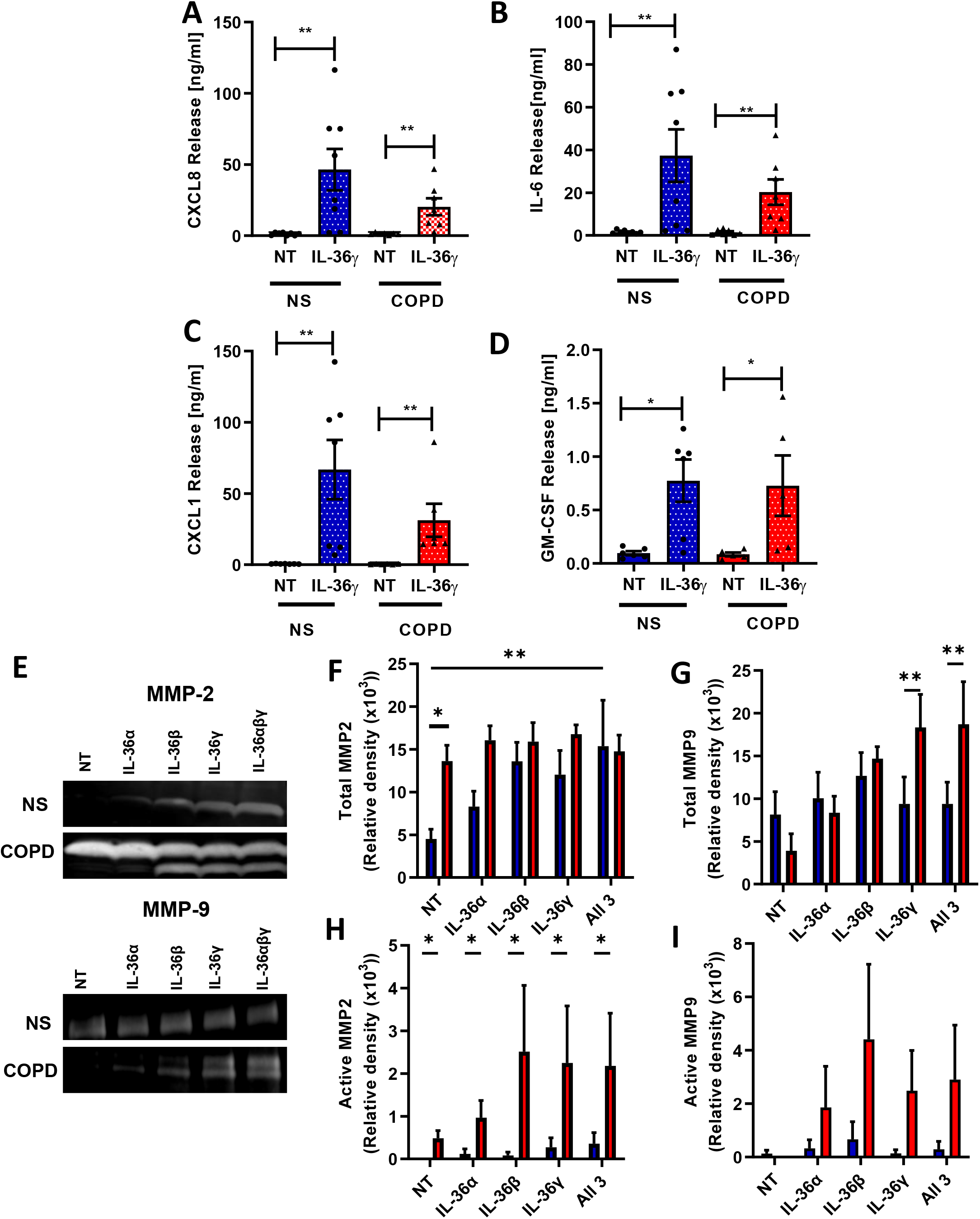
IL-36γ activates small airway fibroblasts leading to chemokine and protease release. Small airway fibroblasts from non-smokers (n=8) and COPD (n=7) patients were cultured in the absence (NT) or presence of 100 ng/ml IL-36γ for 24h. Media was harvested and A) CXCL8, B) IL-6, C) CXCL1 and D) GM-CSF were measured by ELISA. Small airway fibroblasts from non-smokers (n=3) and COPD patients (n=3) were cultured in the absence (NT) or presence of 33 ng/ml IL-36α, IL-36β, IL-36γ or all three in combination and media collected and zymography performed (panel E). Data are means ± SEM and analysed by Kruskal-Wallis test with post-hoc Dunn’s test; * P <0.05, **P<0.01

SAF are found at the site of airways obstruction in COPD and are important in the development of peribronchiolar fibrosis which is the hallmark of early disease (Koo et al., 2018). Matrix metalloproteinases (MMP)-2 and MMP-9 are increased in COPD and are highly activated (Churg et al., 2012; Russell et al., 2002b). SAF from COPD patients release increased levels of active MMP-2 (4.6 fold) and total MMP-9 (4.7 fold) when stimulated with IL-36γ (Fig. 6E), and stimulation of these cells with any of the IL-36 isoforms appears to lead to an increase in activation of MMP-2 (Fig. 6E). Taken together, these data highlight the potential role of SAF in driving neutrophil recruitment and airway remodelling in COPD.

Fibroblast phenotypes are also altered in COPD, with differentiation towards myofibroblasts, leading to greater collagen deposition within the lung (Karvonen et al., 2013). We therefore examined the effect of IL-36γ on these fibrosis markers in comparison with IL-1α, to determine whether other members of the IL-1 family elicited similar effects. *COL1A1* and *COL3A1* gene expression appeared unchanged in SAF from both non-smoker and COPD patients when treated with either IL-36γ or IL-1α (Supplementary Fig. 5A and B). However, *α-SMA*, a myofibroblast differentiation marker was significantly decreased in SAF from both non-smoker and COPD patients by both IL-36γ and IL-1α (Supplementary Fig 5C), suggesting IL-36γ was driving these cells away from a myofibroblast phenotype, to a potentially more proinflammatory and proteolytic state. This was confirmed by an increase in CXCL8, IL-6 and MMP-9 gene expression in these cells (Supplementary Fig 5D-F). Interestingly, as seen previously with protein release, SAEC stimulated with IL-36γ did not respond as robustly as fibroblasts, with only a small but significant change in the gene expression of CXCL8 and IL-6, but no change in MMP2 or MMP9 (Supplementary Fig. 5K-L).

IL-36Ra may be induced by IL-36 cytokines as a negative feedback loop. We therefore also examined the effect of IL-36γ and IL-1α on gene expression of IL-36RN. We observed increased expression of *IL-36RN* in response to IL-36γ in non-smoker SAF, but this was blunted in COPD SAF (Supplementary Fig 5G). Similarly, IL-1α caused a significant increase *IL-36RN* in non-smoker SAF, but this was not seen in COPD SAF (Supplementary Fig 5G). Interestingly, *IL-1RA* was induced in both non-smoker and COPD SAF when stimulated with IL-1α, but when stimulated with IL-36γ the response was again blunted in COPD patients (Supplementary Fig 5H). This suggests that IL-36γ mediated upregulation of IL-36RN is attenuated in COPD, resulting in increased activity of IL-36γ.

### Effect of glucocorticosteroids on IL-36γ responses

As inhaled glucocorticosteroids are commonly used in the treatment of COPD, we examined whether IL-36γ-driven inflammation was steroid-sensitive. SAF from non-smokers and COPD patients were stimulated with IL-36γ and the effect of increasing concentrations of budesonide on the release of the chemokines and cytokines was examined. Interestingly, CXCL8, IL-6 and GM-CSF were all reduced by budesonide in a concentration-dependent manner (Fig. 7A-C). However, CXCL1, one of the major neutrophil recruiting chemokines in COPD, was surprisingly increased by budesonide treatment (Fig. 7D), suggesting that steroid treatment may paradoxically further perpetuate IL-36γ inflammation by increasing neutrophil recruitment.

**Figure 7.**
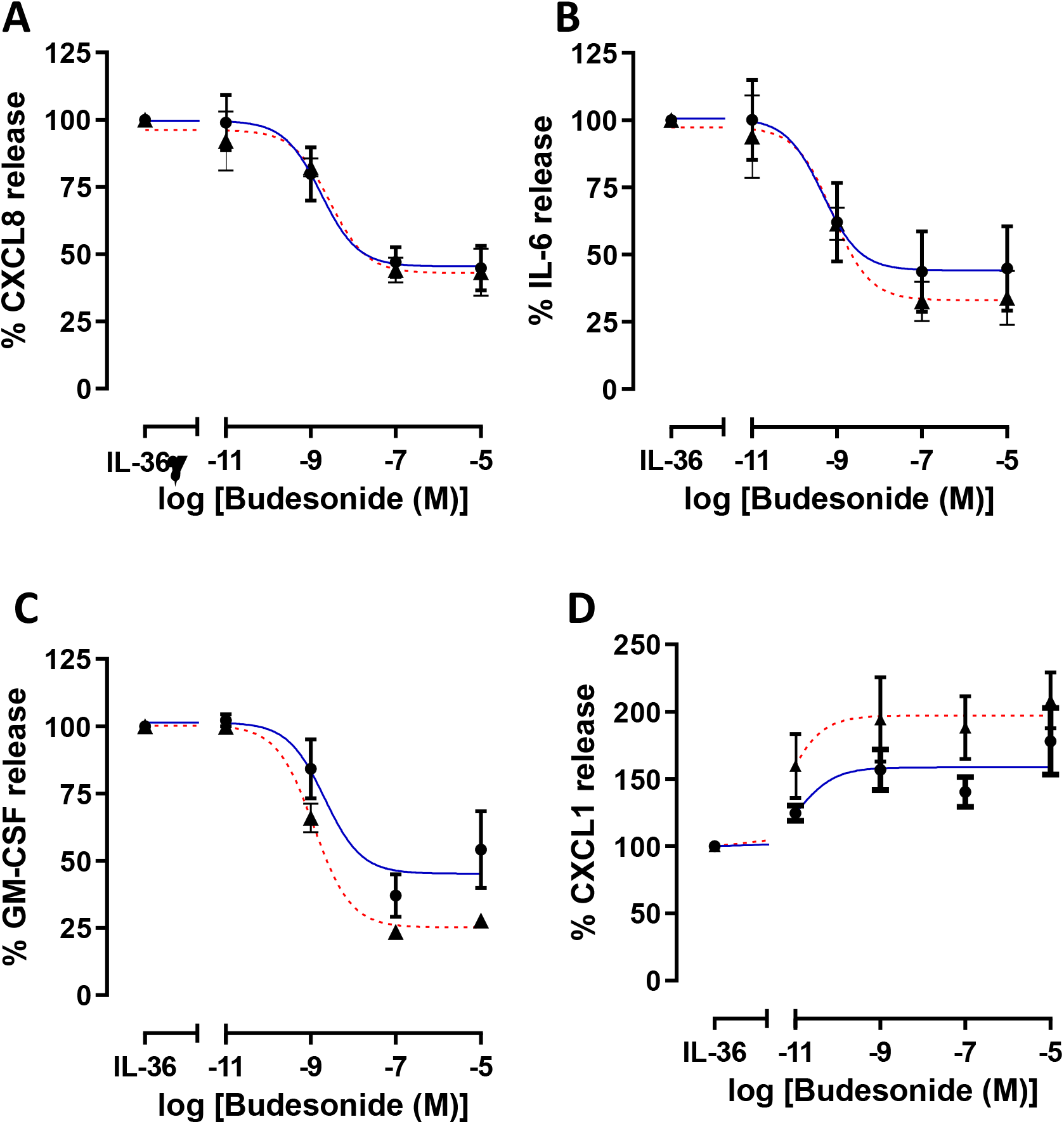
IL-36 stimulation of SAF is glucocorticosteroid sensitive, except the neutrophil chemokine CXCL1 which is induced by budesonide. Small airway fibroblasts from non-smokers (● n=4) or COPD (▲ n=4) were treated with active IL-36γ for 24 hours in the absence or presence of budesonide at varying concentrations, A) CXCL8, B) IL-6, C) GM-CSF and D) CXCL1 release were measured by ELISA. Data are presented as mean ± SEM.

### Serine proteases activate IL-36γ

Previous studies have suggested that intratracheal instillation of IL-36 into the lungs of mice induces neutrophil recruitment (Koss et al., 2021; Ramadas et al., 2012; Ramadas et al., 2011). Recruitment and subsequent activation of neutrophils at the site of inflammation in the lung leads to the release of proteolytic enzymes, such as the serine proteases neutrophil elastase, proteinase-3 and cathepsin G, all of which are elevated in the COPD lung (Stockley, 1999). Elevated numbers of neutrophils and macrophages are found within the lungs of COPD patients (Barnes, 2016b), with both cell types releasing proteases, which when unchecked can lead to damage and remodelling of the lung (Churg and Wright, 2005). Neutrophil elastase, proteinase-3 and cathepsin G have all been suggested to cleave IL-36 cytokines, although there are conflicting results within the literature (Ainscough et al., 2017; Clancy et al., 2018; Henry et al., 2016). We therefore assessed whether neutrophil proteases (neutrophil elastase, proteinase-3 and cathepsin G) or a macrophage protease (MMP-9) could cleave and activate IL-36γ. Utilising a cell free assay, we incubated recombinant full-length IL-36γ with various concentrations of these proteases. Our results show that both cathepsin G and proteinase-3 were capable of cleaving IL-36γ into its active form, whereas neutrophil elastase did not (Fig. 8A), suggesting that neutrophils may activate IL-36γ in COPD.

**Figure 8.**
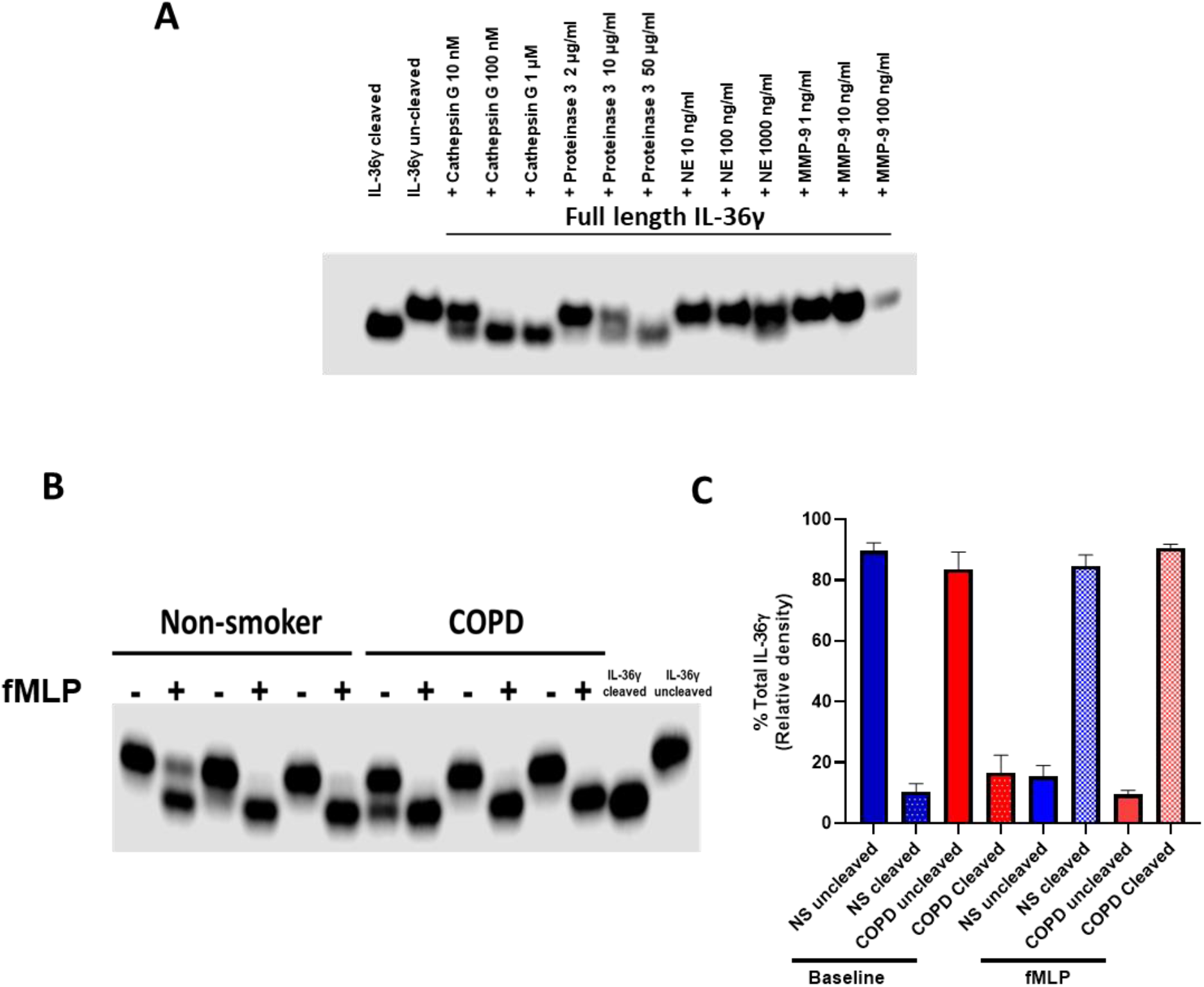
Effect of neutrophil serine proteases on activation of IL-36γ. A) Recombinant full length IL-36γ was incubated with cathepsin G, proteinase 3, neutrophil elastase (NE) or matrix metalloproteinase (MMP)-9 at varying concentrations for 2.5 h and Western blots performed to separate pro- and active (cleaved) IL-36γ. B) Neutrophils from non-smokers (n=6) and COPD patients (n=6) were left at baseline or activated with fMLP and the supernatant collected. Supernatants were incubated with recombinant full length IL-36γ for 2.5h and western blots performed to separate pro- and active (active) IL-36γ (data shown are representative blots). Optical denisty of bands was performed and data are presented as mean ± SEM (panel C).

As it appeared that neutrophil proteases cleaved IL-36γ, we next sought to see whether activated neutrophil products from both non-smokers and COPD patients could cleave and activate IL-36γ. Full-length IL-36γ was converted to the active form by both non-smoker and COPD fMLP-activated neutrophils, although data suggest basally released COPD neutrophil products may be able to cleave IL-36γ (Fig. 8B and C). Elevated numbers of neutrophils within the COPD lung may therefore have the capability to further activate IL-36 cytokines further amplifying the inflammation IL-36γ may cause within the COPD lung.

### Blocking the IL-36R prevents inflammatory cross-talk between COPD SAEC and SAF

As we have shown that IL-36Ra is down-regulated in COPD patients and may exacerbate the inflammatory response of IL-36 cytokines in the COPD lung, we investigated whether re-introduction of IL-36Ra could inhibit IL-36γ driven inflammation. Experiments were devised whereby media from poly I:C stimulated SAEC from COPD patients was transferred to COPD SAF and CXCL1 measured as an output. A schematic of the experiment is depicted in Figure 9A. The transferred media contained IL-36γ (unstimulated: 15.9 pg/ml; stimulated 307.9 pg/ml) and CXCL1 (unstimulated: 1.9ng/ml; stimulated 3.1 ng/ml). Native media from poly I:C stimulated SAEC led to a significant release of CXCL1 from SAF as previously seen, whilst little CXCL1 release was seen when SAF were stimulated with poly I:C alone (Figure 9B). To show that the induction to CXCL1 in the SAF was a consequence of a protein mediator within the media, the media was boiled to denature proteins and SAF treated with this media (Figure 9C). Having established a media transfer system, SAF were then treated for 2 hours with 100 ng/ml of recombinant IL-36Ra, before being treated with SAEC media. Pre-treatment with IL-36Ra led to a significant reduction in CXCL1 release from SAF treated with poly I:C-stimulated SAEC media (Fig. 9D). To confirm these findings, we utilised an IL-36R neutralising antibody. SAF from COPD patients were treated with 50 µg/ml of either isotype control or an antibody which binds the IL-36R and blocks signalling, followed by 100 ng/ ml of IL-36γ. IL-36γ induced CXCL1 release from cells treated with the isotype control, but this was abolished in cells pre-treated with the IL-36R neutralising antibody, showing the antibody inhibited IL-36-mediated CXCL1 release (Fig. 9E). SAF were then treated for 2 hours with the isotype control or IL-36R antibody, before being treated with SAEC media. Pre-treatment with IL-36R led to a significant reduction in CXCL1 release from SAF treated with poly I:C-stimulated SAEC media (Fig. 9F). These data suggest that blocking IL-36R via increasing the reduced endogenous levels of IL-36Ra or by directly blocking the IL-36R using a neutralising antibody, it is possible to prevent the cross talk between virally stimulated COPD SAEC and COPD SAF. These data suggest that blocking IL-36 signalling via these two methods may reduce virally induced IL-36 mediated inflammation in COPD.

**Figure 9.**
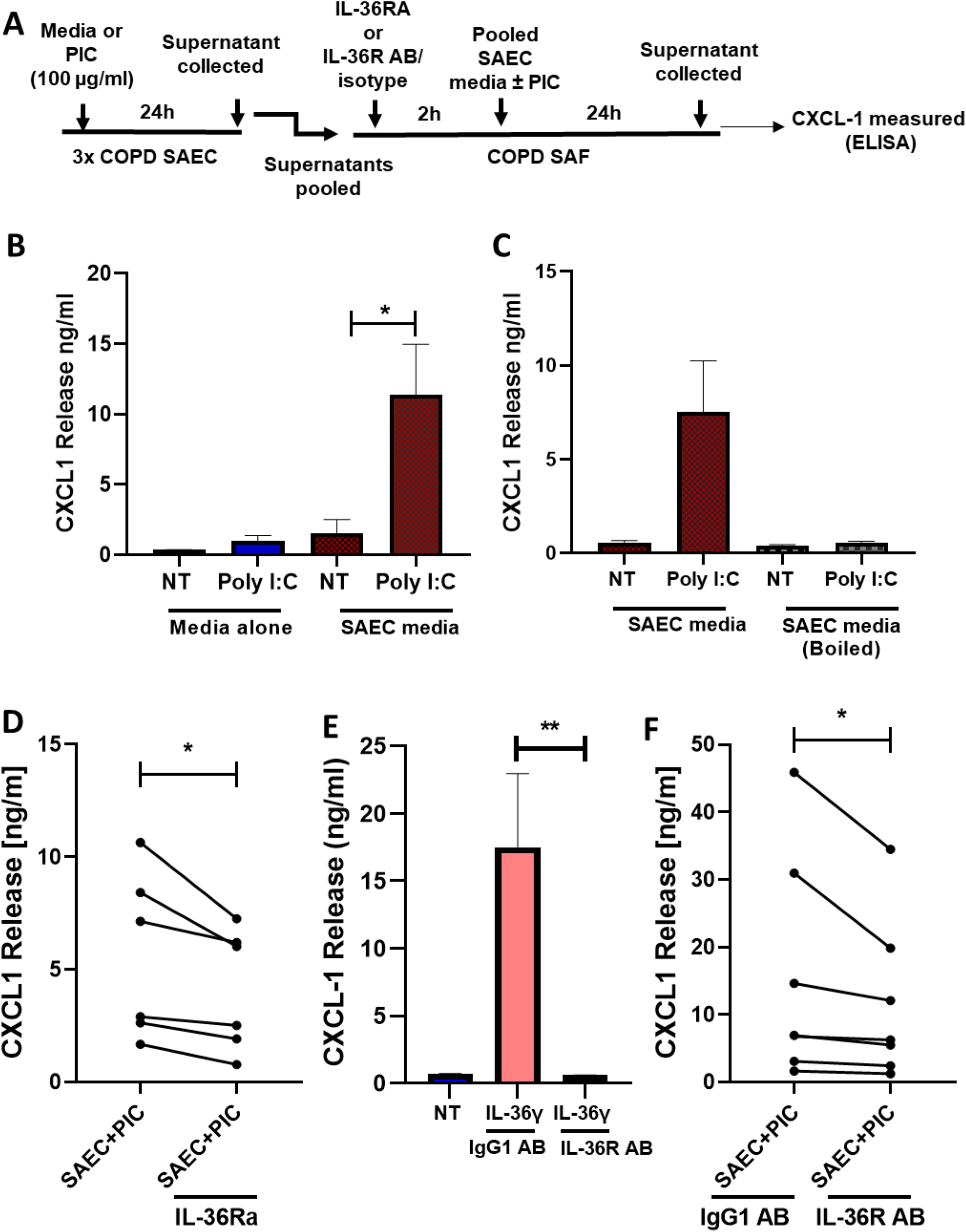
Optimisation of therapeutic IL-36R inhibition experiments. A) Schematic of experimental procedure. B) Small airway fibroblast (n=6) were treated with diluted media, media + Poly I:C, media alone treated SAEC or Poly I:C treated SAEC media (ranging from 10-200 fold) for 24 hours(pooled media from 3 COPD patients). CXCL1 levels were then measured. C) Small airway fibroblast (n=4) were treated with diluted (ranging from 10-200 fold) media alone treated SAEC or Poly I:C treated SAEC that had and hadn’t been boiled for 24 hours. CXCL1 levels were then measured. D) Small airway fibroblast (n=6) were treated with diluted (ranging from 10-200 fold) media alone treated SAEC or Poly I:C treated SAEC with or without pre-treatment for 2 hours with recombinant IL-36Ra (100 ng/ml). CXCL-1 levels were then measured. E) Small airway fibroblast (n=4) were pre-treated for 2 hours with isotype control (IgG1) antibody or a IL-36R neutralising antibody and then treated with 100 ng/ml of IL-36γ for 24 hours. F) Small airway fibroblast (n=7) were treated with diluted (ranging from 10-200 fold) media alone treated SAEC or Poly I:C treated SAEC with pre-treatment with either isotype control (IgG1) antibody or a IL-36R neutralising antibody for 2 hours. CXCL-1 levels were then measured. Data are presented as mean ± SEM analysed by either Kruskal-Wallis with post hoc Dunns or Wilcoxon matched-pairs signed rank test; ** P <0.01, * P <0.05.

## Discussion

Neutrophilic inflammation is characteristic of COPD airways, and this is associated with increased expression of neutrophil chemoattractants, CXCL1 and CXCL8 (Keatings et al., 1996; Traves et al., 2002). Our study describes a novel mechanism for the amplification and perpetuation of chronic neutrophilic inflammation in COPD patients. We confirm previous findings that IL-36γ, but in this study not IL-36α, is elevated in the lungs of COPD patients but show for the first time that there are reduced levels of the endogenous IL-36 receptor inhibitor, IL-36Ra, suggesting enhanced IL-36 signalling in COPD lungs. Examining multiple cell types from non-smokers, smokers and COPD patients we established that elevated levels of IL-36γ in COPD BAL and nasal fluid is potentially derived from epithelial cells, which release higher basal IL-36γ levels than cells from non-smokers. Examining different cell types, we show that SAF appear to be the main IL-36γ effector cell, releasing marked amounts of chemokines, proinflammatory cytokines and proteases upon stimulation. These elevated chemokines recruit neutrophils into the lung that may activate the elevated IL-36γ, by releasing serine proteases capable of cleaving IL-36γ into its active form; our data suggest these proteases are cathepsin G and proteinase-3, but not neutrophil elastase. We show that treating SAF with IL-36Ra or by using a therapeutic antibody which blocks IL-36R mediated signalling, inhibits viral-induced IL-36-mediated inflammatory cross-talk between SAEC and SAF in COPD. IL-36γ may therefore drive COPD pathophysiology via the recruitment and activation of neutrophils into the lung, leading to small airway remodelling, emphysema, and mucus hypersecretion (Fig. 10).

**Figure 10.**
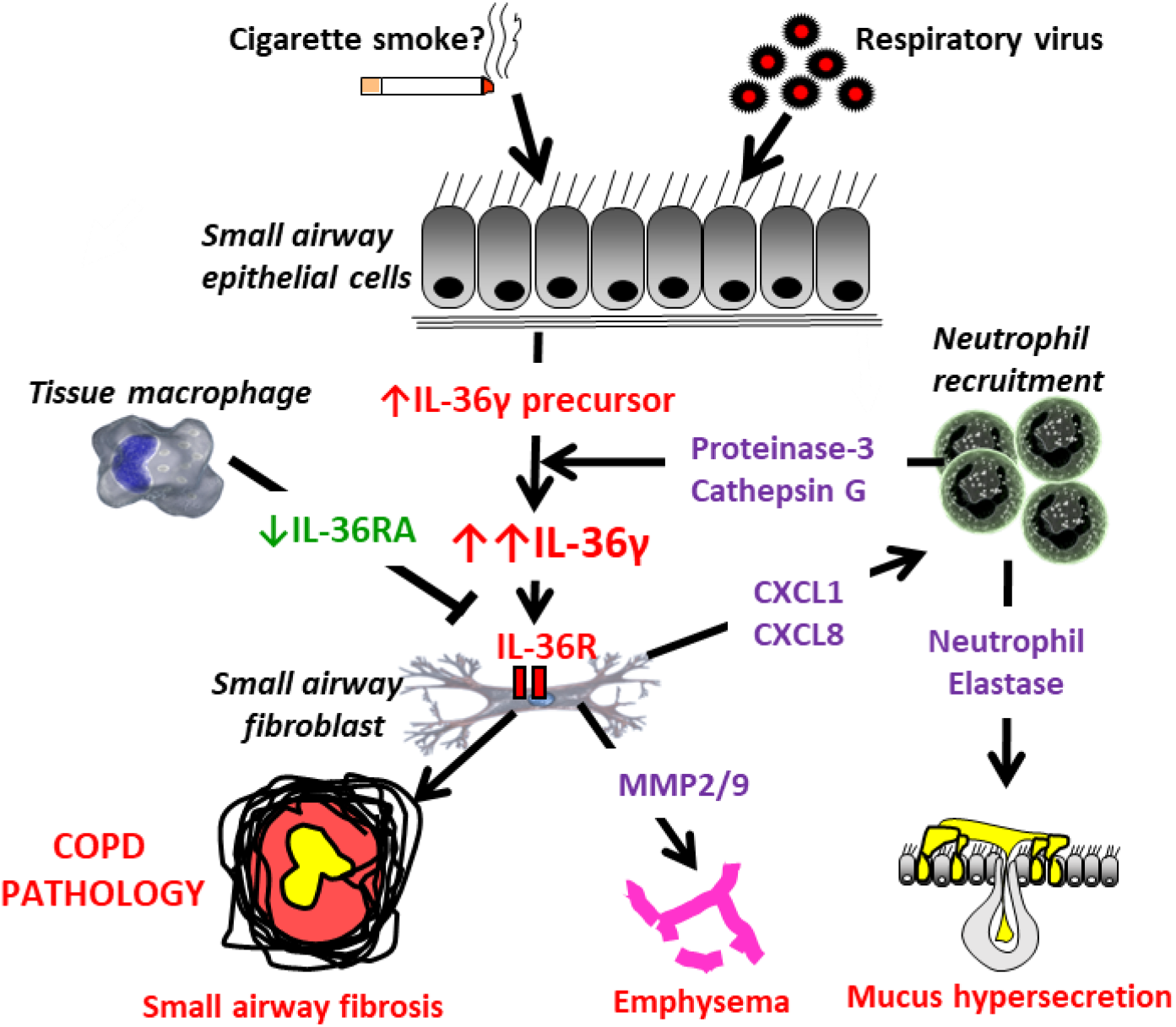
Schematic outlining the role of IL-36γ in COPD pathophysiology. Small airway epithelial cells are a major source of IL-36γ in the COPD lung and the release of IL-36γ is elevated at baseline in these patients. These elevated levels can be exacerbated by viral infection, which may be perpetuated in those who smoke. The resultant pro-IL-36γ is cleaved by neutrophil derived proteases such as cathepsin G and proteinase 3, generating the active form of IL-36γ. This acts on small airway fibroblasts leading the release of MMP-2 and MMP-9 as well as expression of the chemokines CXCL-1 and CXCL-8. These recruit neutrophils and perpetuate the cycle of neutrophilic inflammation. This process is amplified in COPD patients due to the loss of the endogenous IL-36 receptor, IL-36Ra, from macropahges. The released elastases and MMPs contribute to all three pathophysiological features of COPD, small airway remodelling, emphysema and mucous hypersecretion.

IL-36γ was the only IL-36 family agonist detected in BAL fluid and levels were elevated in both smokers and COPD patients. Previously, Kovach *et al*., reported that IL-36α may also be elevated in the BAL of COPD patients and smokers, as well as IL-36γ, and in agreement with our study, that IL-36β was undetectable. These data suggested this may be due to smoking (Kovach et al., 2020). However, exposure of SAEC to cigarette smoke extract *in-vitro*, did not stimulate release of IL-36γ in our study. This again contrasts with Kovach *et al*., who found increased release of IL-36γ in response to CSE. However, these cells were also of bronchial origin and may reflect a different response to CSE to epithelial cells for peripheral airways, which are believed to be the major effector cell in COPD. This may also reflect differences in chronic exposure as opposed to a more acute exposure in our *in-vitro* cell system. Others have reported increased *IL-36γ* mRNA expression in response to CSE in bronchial epithelial cells, but only after 8 hours and was reduced 2-fold after 24 hours stimulation (Parsanejad et al., 2008). Although smoking may affect IL-36γ release it appears that this unlikely to alone account for the elevated basal release of COPD SAEC.

A clear difference between smokers and COPD patients was reduced expression of IL-36Ra. The loss of this natural antagonist of the IL-36R may amplify and perpetuate IL-36 mediated inflammation and cause greater effects within the lungs of COPD patients compared to those who smoke. This is pertinent as we were unable to detect IL-38, another antagonist of the IL-36R. This may be very relevant in COPD as IL-36Ra binds to the IL-36R with a greater affinity and for greater duration than IL-36 agonists, suggesting its loss may be detrimental in overcoming IL-36 agonist mediated inflammation (Zhou et al., 2018). This is further evident as the loss of IL-36Ra is prevalent in GPP, where missense mutation in the *IL-36RN* gene leads to IL-36Ra deficiency and drives this disease (Marrakchi et al., 2011). There are no indications to date of a similar genetic defect in COPD, however clinical trials are underway in GPP, assessing whether targeting the IL-36R with a blocking monoclonal antibody (spesolimab) prevents and reverses this disease (Bachelez et al., 2019). Our data suggest that elevated levels of the IL-36 receptor agonist, IL-36γ, in COPD are likely to be greatly amplified by the reduced secretion of IL-36Ra and that blocking IL-36R may be of clinical benefit.

IL-36 cytokine levels were also examined via nasosorption. This non-invasive technique can give insight into the cytokine profile found within the lung, with a strong correlation between the levels of inflammatory cytokines in the nasal mucosa and the lung (Batista et al., 2016; Thwaites et al., 2018). This technique, as well as being non-invasive also has the potential advantage of being more sensitive in detecting cytokine levels compared to BAL fluid, as we see much higher concentrations of cytokines can be detected (ng/ml compared to pg/ml in BAL fluid), as these are not diluted by the additional of saline used for the procurement of the BAL fluid. We show a significant increase in the levels of IL-36γ in these samples, whilst showing no changes in the levels of either IL-36α and IL-36β further confirming our findings from the BAL samples.

We next sought to identify the cellular source of IL-36γ within the lung. In psoriasis, keratinocytes are the major cell type that releases IL-36 cytokines, but the cellular source in the lung has not been established. Interestingly, IL-1 is highly abundant in all cell types, but the IL-36 related cytokines appear to be more cell specific, with an expression profile suggesting induction in epithelial cells (Wang et al., 2019). When examining the source of IL-36γ in the lung we could detect IL-36γ release only from SAEC, with levels being undetectable in SAF and TMφ, suggest the epithelium as a major source. Previous data has also suggested that bronchial epithelial cells release IL-36 cytokines in response to ds RNA. Our data using poly(I:C) confirm these findings, and again showed no released from TMφ and SAF with this stimulus.

The epithelium is the major site of viral infection within the lung (Vareille et al., 2011) and our data suggest that IL-36γ can be further induced in COPD SAEC when treated with the viral mimetic poly (I:C). Upper respiratory tract viral infections are a major cause of COPD exacerbations and a leading cause for hospitalization for COPD patients (Wedzicha and Donaldson, 2003). Lung function declines during an exacerbation, and may never completely recover, showing the contribution of these episodes to disease progression, in addition to the high overall costs of hospitalization. These data suggest that IL-36γ could be a potential biomarker for viral infections in COPD, especially as IL-36 cytokines can be detected in nasal samples and therefore patients can be non-invasively tested during these episodes. Acute exacerbations of COPD are also triggered by bacterial infections, particularly *Haemophilus influenzae*, but treatment of SAEC with *H. influenzae*, had no effect on the release of IL-36γ; suggesting that blocking IL-36 would be more effective in preventing viral exacerbations.

Identification of the main IL-36 effector cell in the lung is crucial in understanding the role of the elevated levels of IL-36γ in COPD. We found that TMφ released very little cytokines in response to IL-36γ, suggesting these to not be a major effector cell in the lung. TMφ from smokers and COPD patients were also not stimulated by any IL-36 cytokines to release CXCL8 (Supplementary Fig 4C). Previous studies have suggested that monocyte-derived M2, but not M1, macrophages release IL-6 in response to IL-36β (Dietrich et al., 2016). This is discrepant with the data presented here and may reflect differences between tissue resident cells and monocyte-derived macrophages, although the aged human lung is thought to consist mainly of monocyte-derived macrophages (Byrne et al., 2020). Our data agrees with others who showed that, bronchial epithelial cells stimulated with IL-36γ released CXCL8 and IL-6 (Zhang et al., 2017). In contrast, SAF secreted much higher concentrations of cytokines, chemokines and active MMPs in response to IL-36 cytokines, suggesting that these are likely to be the major effector cell in the lung. Lung fibroblasts, along with colonic fibroblasts, have previously been shown to secrete cytokines, chemokines and MMPs in response to IL-36 (Chustz et al., 2011; Scheibe et al., 2019). Epithelial cells grown at air liquid interface have been shown to release IL-36 cytokines from the basolateral surface and therefore fibroblasts being the main effector cell correlates with this finding (Chustz et al., 2011). IL-36 stimulation also induced extracellular matrix deposition from human fibroblasts (Sun et al., 2013), however we found no change in the gene expression of collagen genes and a down-regulation of the myofibroblast marker α-SMA in response to IL-36, suggesting a change in fibroblast phenotype towards a more remodelling than profibrotic type of cell with increased expression of MMP rather than a matrix generating cell.

Systemic glucocorticosteroids are commonly used in the treatment of COPD exacerbations in an attempt to reduce the increased inflammation associated with these infective episodes. We therefore tested whether IL-36γ-mediated inflammation was steroid-sensitive. Interestingly, CXCL8, IL-6 and GM-CSF induction by IL-36γ in SAF was steroid-sensitive, with release reduced by ∼50-60% by high concentrations of budesonide. However, a major neutrophil chemokine, CXCL1, was paradoxically induced by budesonide in the presence of IL-36γ, suggesting that administration of steroids to patients with elevated IL-36γ may be pro-inflammatory via increased CXCL1 and increased neutrophil recruitment. Previous data have suggested that CXCL1 levels are unaffected by inhaled glucocorticosteroids in COPD patients, in contrast to CXCL8 (Inui et al., 2018). *In vitro* studies are conflicting, with studies suggesting both steroid sensitivity and insensitivity to the same and different stimuli (Lo et al., 2014; Shieh et al., 2014). Nevertheless, our data suggest that giving a steroid during an exacerbation when IL-36γ may be induced via a virus, could lead to further neutrophil recruitment and thus be detrimental to the patient.

SAF release high levels of the two main neutrophil chemokines, CXCL1 and CXCL8, in response to IL-36γ. These chemokines recruit neutrophils from the blood into the lungs by binding to the common chemokine receptor CXCR2. Higher numbers of COPD neutrophils migrate to CXCL1 compared to cells from non-smokers, and this cytokine is markedly elevated in the lungs of COPD patients (Dunne et al., 2019; Traves et al., 2002). To enter the lung these neutrophils must migrate though the tissue releasing proteases as they travel. In COPD this migration is altered with an increase in speed that is less directional (Sapey et al., 2011) and this can lead to excess tissue damage. Therefore, the elevation of both CXCL1 and CXCL8 can increase the recruitment of neutrophils from the blood into the COPD lung and in doing so cause excessive tissue damage due to their dysregulated migratory path.

Once in the lung, neutrophils degranulate and release several proteases including the serine proteases neutrophil elastase, cathepsin G and proteinase-3. Here, we show that cathepsin G and proteinase-3 cleave IL-36γ into its active form, which has been shown to have a 500-fold greater activity than the uncleaved secreted cytokine. These data contrast with some studies that show that neutrophil elastase is the main activator of IL-36γ (Henry et al., 2016) but this is also controversial with others failing to show this response (Ainscough et al., 2017; Bassoy et al., 2018). However, as neutrophils are increased and activated in COPD it is plausible that they activate IL-36γ leading to marked increases in the potency of IL-36γ within the COPD lung, thus amplifying and perpetuating neutrophilic inflammation.

Finally, we showed that blocking the IL-36R by treating SAF with either recombinant IL-36Ra or an IL-36R blocking antibody reduced CXCL1 release from these cells when stimulated with media from poly I:C stimulated COPD SAEC. These data suggest that re-introducing the natural inhibitor of the IL-36R or directly blocking the receptor with a therapeutic antibody, such as spesolimab, may reduce viral induced inflammation via the IL-36 pathway. We have recently shown in an *in-vivo* model of a COPD viral exacerbations, knockout of the IL-36R reduces lung inflammation and neutrophil recruitment (Koss et al., 2021). Our cross-talk experiments shows that attenuating the activity of the IL-36R in primary human cells from COPD patients may also lead to reduced inflammation, suggesting that this may be translated into humans and a potential therapeutic option.

The regulation of IL-36 cytokines is complex, with the requirement for extracellular protease activation and their modulation by the antagonistic IL-36Ra and IL-38. These different IL-36 family members appear to be released from different cell types, suggesting a complex interactive cell network (Bassoy et al., 2018). The effects of IL-36 cytokines are similar to those of related IL-1 cytokines, which are released predominantly via a different mechanism, which involves intracellular activation via the inflammasome (Martinon et al., 2002). It is possible that IL-1 cytokines provide the initial inflammatory process in host defence and that IL-36 cytokines are activated with greater stimulatory triggers or more prolonged stimulation, resulting in greatly amplified and persistent neutrophilic inflammation, as found in COPD patients.

Overall, our data suggest IL-36γ is elevated in COPD, and is released predominantly by epithelial cells, leading to the activation of fibroblasts, inducing neutrophilic recruitment which further activates IL-36γ, inducing protease release and inflammation, all of which drive with COPD pathophysiology (Fig. 10). This mechanism is amplified by a reduction in IL-36Ra from macrophages and by the marked activation of IL-36γ by serine proteases released from the activated recruited neutrophils. We suggest that this is a major mechanism for amplification of lung inflammation in COPD leading to persistent inflammation and disease progression. We show that the blocking of the IL-36R by either increasing the endogenous levels of IL-36 This suggests that targeting IL-36 cytokines, for example with neutralizing antibodies against the receptor or addition of IL-36Ra, is a promising new therapeutic opportunity for the treatment of COPD.

## Material and methods

### Reagents

Recombinant IL-1α, IL-36α, IL-36β, IL-36γ, IL-36RA, TNF-α, neutrophil elastase, cathepsin G and proteinase 3, and anti-IL-36γ antibody (AF2320) were purchased from R&D Systems (Abingdon, UK). Rabbit Anti-Goat Immunoglobulins/HRP (P0449) was purchased from Agilent (Santa Clara, USA) Poly I:C was purchased from Sigma-Aldrich (Poole, UK). Non-typeable *H. influenzae* were obtained from the National Collection of Type Cultures (strain: 1269) and were heat-killed by incubation at 65 °C for 10 min as described previously (Taylor et al., 2010). Cigarette smoke extract (CSE) was generated as previously described (Yanagisawa et al., 2017) from full-strength Marlboro cigarette (Phillip Morris, London, UK). Budesonide was purchased from Fisher Scientific UK Ltd (Loughborough, UK).

### Bronchoalveolar Lavage

Bronchoalveolar lavage was performed as described previously (Culpitt et al., 2003; Russell et al., 2002a). Briefly, BAL was collected from the right middle lobe by instilling 60 ml of warmed 0.9% (wt/vol) normal saline into the lung to a maximum of 240 ml. BAL was filtered and centrifuged to remove cells and stored at –80°C. See (Supplementary table 1) for patient demographics.

### Induced sputum

Induced sputum was collected and processed following a modification of Pin et al., (Pin et al., 1992) as reported previously (Costa et al., 2008). Briefly, subjects inhaled increasing concentrations of hypertonic saline solution (3%, 4% and 5% [weight/volume]) for 7 min at each concentration. The opaque, gelatinous portions of sputum were selected and centrifuged at 300g. The viscous sample was weighed and treated dithiothreitol (DTT) diluted to 0.1% (weight/volume) with distilled water. The sample was solubilised by vortexing. Four volumes of Dulbecco phosphate-buffered saline solution were added to the sample to give a final concentration of 0.05% (weight/volume) DTT. The supernatant was collected and stored at − 80°C (Costa et al., 2008). See (Supplementary table 2) for patient demographics.

### Nasosorption

Nasosorption was performed using Nasosorption™ FX·I device (Hunt Developments UK Ltd) as described by others (Habibi et al., 2020; Jha et al., 2021; Morton et al., 2021; Thwaites et al., 2018).Briefly, a Nasosorption™ FX·I device containing a synthetic absorptive matrix (SAM) was inserted into each nostril for 60 seconds for sample collection. Each SAM was detached and placed into 300 µl of elution buffer and vortexed for 30 seconds and transferred to a Costar SPIN-X bucket (Sigma Millipore). Sample is then centrifuged for 20 min at 16,000 x g in a mini-centrifuge cooled to 4 °C. Supernatant is then stored at − 80°C. See (Supplementary table 3) for patient demographics.

### Primary human lung cells

Lung tissue macrophages were isolated from lung parenchyma tissue as described previously (Belchamber et al., 2019). Lung tissue was assessed as being non-cancerous and obtained from samples during tissue resection for lung cancer or emphysema. The subjects were matched for age and smoking history (Supplementary Table 4) Human primary small airway epithelial cells (SAECs) were cultured as previously described (Baker et al., 2016). The subjects were matched for age and smoking history (Supplementary Table 5). Small airway fibroblasts were cultured by micro-dissecting out 5 small airways from lung parenchymal tissue and growing via an outgrowth method (Supplementary Table 6). Lung homogenate samples were obtained from an established tissue bank linked to an established patient registry which has previously been used (Ding et al., 2004) (Supplementary Table 7). COPD patients had significantly worse lung function compared to controls. Subjects provided informed consent, and the study was approved by the NRES London-Chelsea Research Ethics committee (study 09/H0801/85).

### Real time PCR

Total RNA was extracted from cells and reverse-transcribed, as described previously (Baker et al., 2016). Gene expression was determined by Taqman real-time PCR on a 7500 Real Time PCR system (Applied Biosystems, Life Technologies Ltd, Paisley, UK) using the assays IL-36α (Hs00205367), IL-36β (Hs00758166), IL-36γ (Hs00219742), IL-36RA (Hs01104220), CXCL8 (Hs00174103), IL-6 (Hs00174131), COL1A1 (Hs00164004), COL3A1 (Hs00943809), α-SMA (Hs05032285) and MMP-9 (Hs00957562). GNB2L1 (Hs00272002) gene expression was used as the housekeeping gene and data presented as δδCT relative to baseline.

### Zymography

MMP2 and MMP9 enzyme activity were measured by zymography using Novex Zymogram Gelatin Gels (Thermo Fisher Scientific). Fibroblast supernatant were diluted in Novex Tris-Glycine SDS sample buffer (Thermo Fisher Scientific) and ran on zymogram gel. After electrophoresis, gels were incubated with Novex zymogram renaturing buffer (Thermo Fisher Scientific) and incubated in Novex zymogram developing buffer (Thermo Fisher Scientific) for 18 h at 37°C. After incubation, gels were stained with a Colloidal Blue Staining Kit (Thermo Fisher Scientific) and imaged.

### ELISA

CXCL8, CXCL1, IL-6, GM-CSF, IL-36α and IL-36β were quantified using commercially available ELISA kits (R&D Systems), according to the manufacturer’s instructions; The lower limit of detection for these assays were 31.2 pg/ml (CXCL1/8, IL-6), 15.6 pg/ml (GM-CSF) and 12.5 pg/ml (IL-36α/β). IL-36γ and IL-36RA were quantified using commercially available ELISA kits (AdipoGen life sciences, Epalinges, Switzerland); The lower limit of detection for these assays were 3.9 pg/ml (IL-36γ) and 0.5 ng/ml (IL-36RA).

### IL-36γ cleavage

Neutrophils were isolated from whole blood from both non-smokers and COPD patients using dextran red blood cell sedimentation, followed by centrifugation using a discontinuous Percoll gradient. Neutrophils were collected from the 81% (v/v) / 67% (v/v) interface, washed in phosphate buffered saline, cells were re-suspended at 10^6^ cells/ml in Reaction Buffer (50 mM HEPES, (Ph7.5) 75mM NaCl, 0.1% (w/v) CHAPS). Cells were treated with 10μM cytochalasin prior to stimulation with 10μM fMLP for 1h (or 0.1% (v/v) DMSO vehicle control) at 37°C. Cells were then centrifuged, and the supernatant removed and stored at -80°C. IL-36γ (uncleaved-1F9) at 1000ng/ml was then incubated at 37 °C for 2.5 h with non-smoker and COPD neutrophil supernatants, Cathespsin G (10,100,1000 nM) (Sigma Millipore), Proteinase 3 (2,10& 50 μg/ml) (Sigma Millipore), neutrophil elastase (10,100 &1000 ng/ml) (Sigma Millipore) or MMP-9 at (1,10 & 100 ng/ml) (R&D Systems Europe) in protease buffer (50mM HEPES (pH 7.5) 75mM NaCl, 0.1% CHAPS). 10 μl (100 ng) of reaction buffer was removed, boiled with Lamaelli buffer for 5 min before loading onto a 4-12% gel using MES buffer. Proteins were transferred to a nitrocellulose membrane before probing using a primary anti IL-36γ antibody (Biotechne (R&D) UK) overnight at 4°C and a secondary goat antibody (Dako) for 1 hour at room temperature to visualise any protein cleavage.

### IL-36RA and IL-36R antibody treatment experiments

SAEC from 3 COPD patients were treated with Poly I:C (100μg/ml) for 24 hours to induce IL-36γ and supernatants collected and pooled. For IL-36RA experiments, pooled supernatant was transferred to COPD SAF and incubated for 24 hours, controls of media alone and or media containing Poly I:C (100 µg/ml) were used. SAF cells were then treated for 2 hours with 100 ng/ml of recombinant IL-36RA, before being treated with SAEC media. For IL-36R antibody experiments, SAF were preteated for 2 hours with either IgG1 isotype control or human anti-IL-36R Ab (Boehringer Ingelheim) 50 μg/ml in sterile PBS, before being treated with SAEC media. In these experiments a maximum of 340 pg/ml of CXCL1 (could have been transferred in the SAEC media to the SAFs, therefore release above this level was because of mediators within the media. To show that the induction to CXCL1 in the SAF was a consequence of a mediator within the media, the media was boiled for 5 mins at 100 degrees and SAF treated with this media and CXCL1 release measured.

### Statistical analysis

Data are expressed as means ± sem. Results were analyzed by using Mann-Whitney tests, paired or nonpaired Student’s t tests, 1- or 2-way ANOVA and Kruskal-Wallis for repeated measures with Dunn’s or Bonferroni post-tests. GraphPad Prism 9 software (GraphPad Software, La Jolla, CA, USA) was used for analyses. Values of P ≤ 0.05 were considered statistically significant.

## Supporting information

All supplementary figures

## Acknowledgments

Special thanks to Professor Jim Hogg (University of British Columbia, Canada) for kindly providing peripheral lung tissue samples and the nurses at the NIHR Respiratory Disease Biomedical Research Unit at the Royal Brompton and Harefield NHS Foundation Trust and Imperial College London.

## Author contributions

JRB was involved in the design, implementation of the experiments and the writing of the manuscript. PF and HO were involved in the implementation of the experiments as well as providing technical expertise. CK, KE-K, MT, JF, SE, PJB and LD were involved in experimental design, interpretation of data, as well as the reviewing of the manuscript. All authors contributed to scientific discussions, revision of the manuscript and had full access to all the data and agreed to submit for publication.

