## Supplementary material for "Imbalance between IL-36 receptor agonist and antagonist drives neutrophilic inflammation in COPD": All supplementary figures

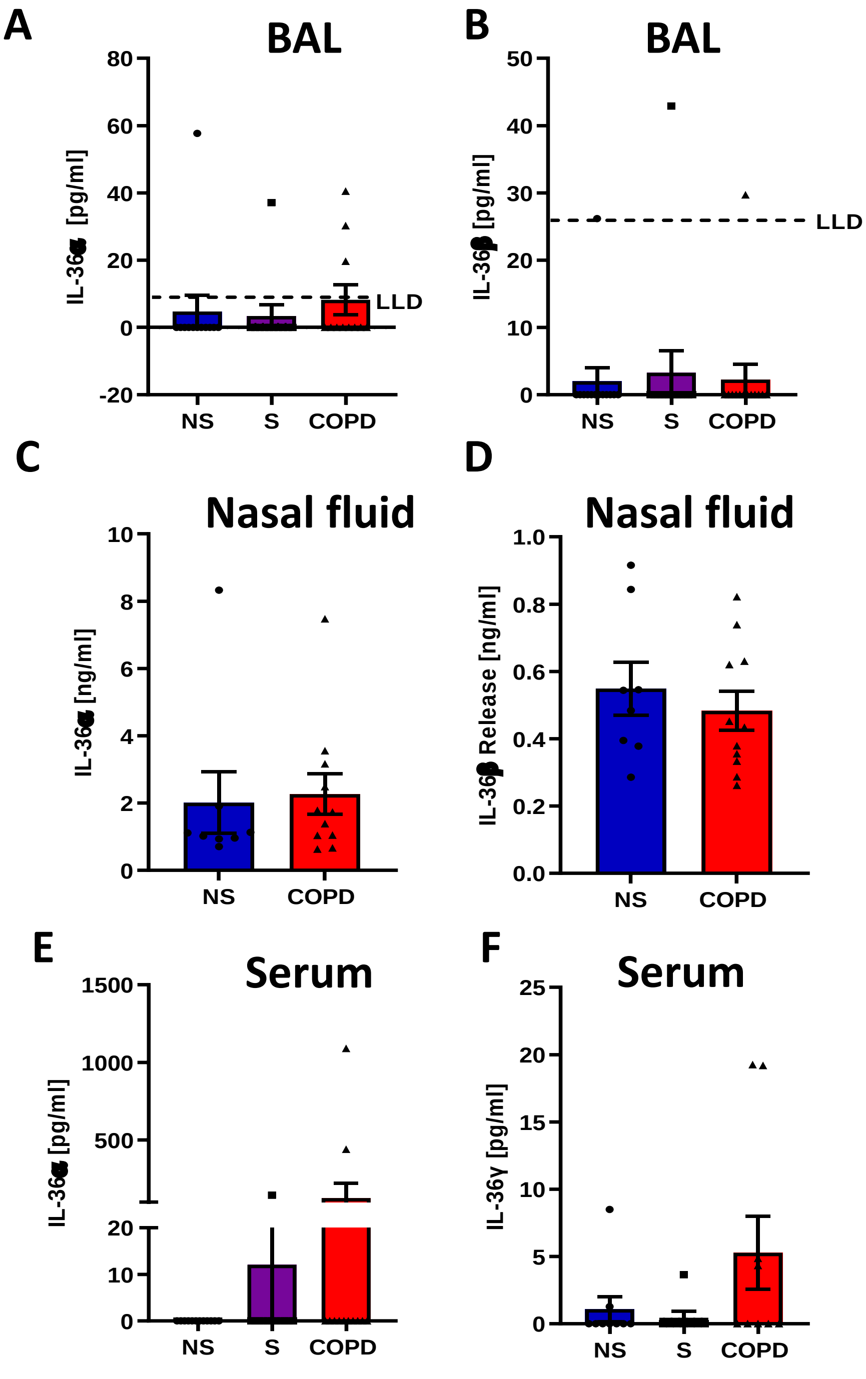


**Supplementary Figure 1. IL-36 cytokines levels in human samples.**

A) IL-36α and B) IL-36β were measured in bronchoalveolar lavage fluid of non-smokers (n=12), smokers (n=11) and COPD (n=11) patients by ELISA. C) IL-36α and D) IL-36β were measured in nasal lining fluid of non-smoker (n=8) and COPD (n=11) patients by ELISA. E) IL-36α and F) IL-36γ were measured in serum samples of non-smokers (n=11) smokers (n=12) and COPD (n=12) patients by ELISA. Data are mean ± SEM and LLD = lower limit of detection.

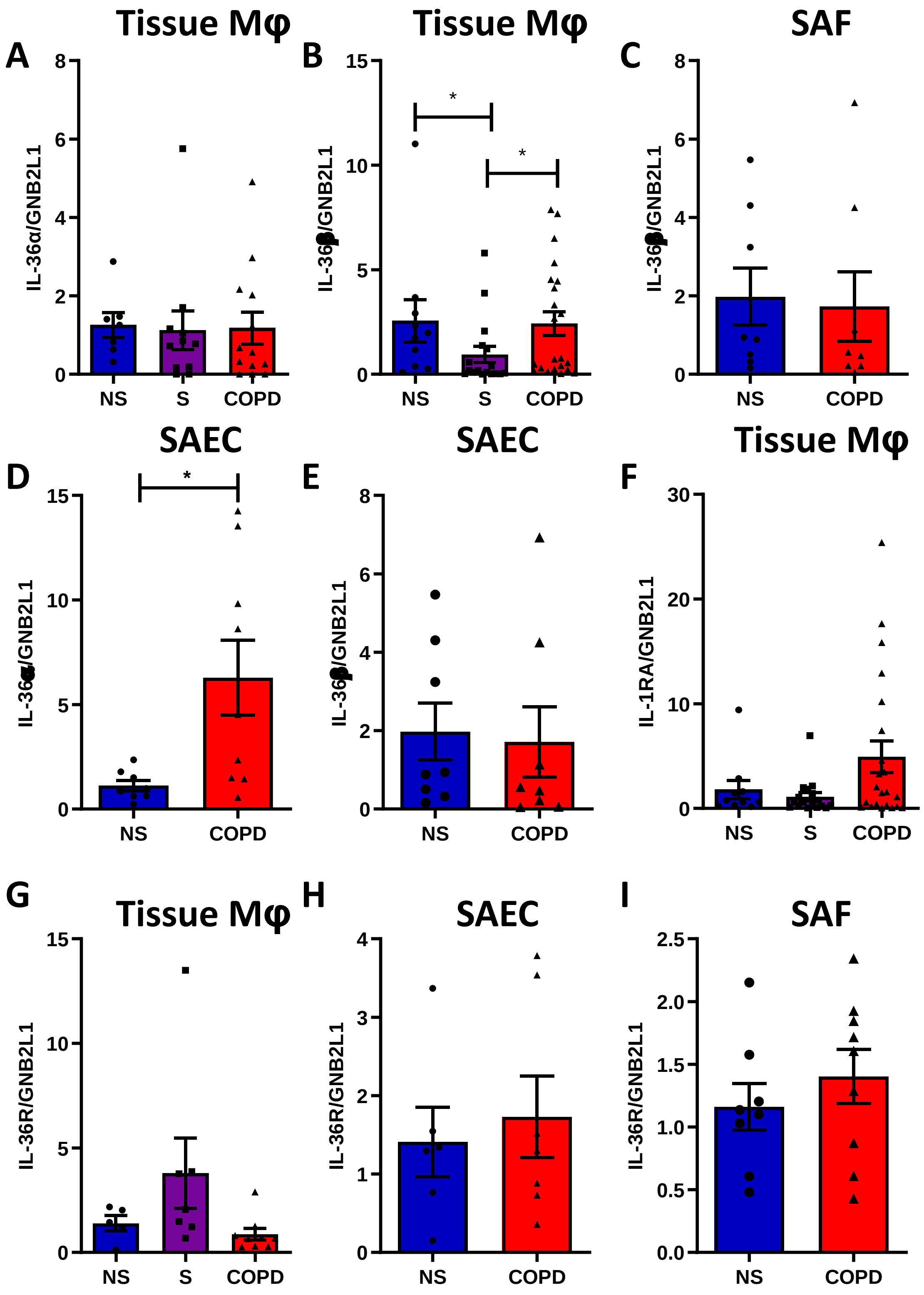

A
Tissue Mφ
IL-36α/GNB2L1
NS
S
COPD
B
Tissue Mφ
IL-36β/GNB2L1
*
*
C
SAF
IL-36β/GNB2L1
D
SAEC
IL-36α/GNB2L1
*
E
SAEC
IL-36β/GNB2L1
F
Tissue Mφ
IL-1RA/GNB2L1
G
Tissue Mφ
IL-36R/GNB2L1
H
SAEC
IL-36R/GNB2L1
I
SAF
IL-36R/GNB2L1

**Supplementary Figure 2. Expression of IL-36 isoforms and receptor in human lung tissue macrophages, small airway fibroblasts and small airway epithelial cells.** A) IL-36α and B) IL-36β gene expression were measured in lung tissue macrophages from non-smokers (NS) (n=7-11), smokers (S) (n=11-17) and COPD patients (n=13-22). C) IL-36β gene expression was measured in SAF from non-smokers (n=8) and COPD patients (n=8). Gene expression of D) IL-36α and E) IL-36β were measured in small airway epithelial cells from non-smokers (n=8) and COPD patients (n=9). F) IL-1RA gene expression was detected in non-smoker (n=10), smoker (n=17) and COPD (n=22) tissue macrophages. IL-36 receptor gene expression was measured in G) tissue macrophages H) SAEC and I) SAF. Data are means ± SEM and analysed by Kruskal-Wallis test with post-hoc Dunn's test or by Mann-Whitney U test; * $P < 0.05$.

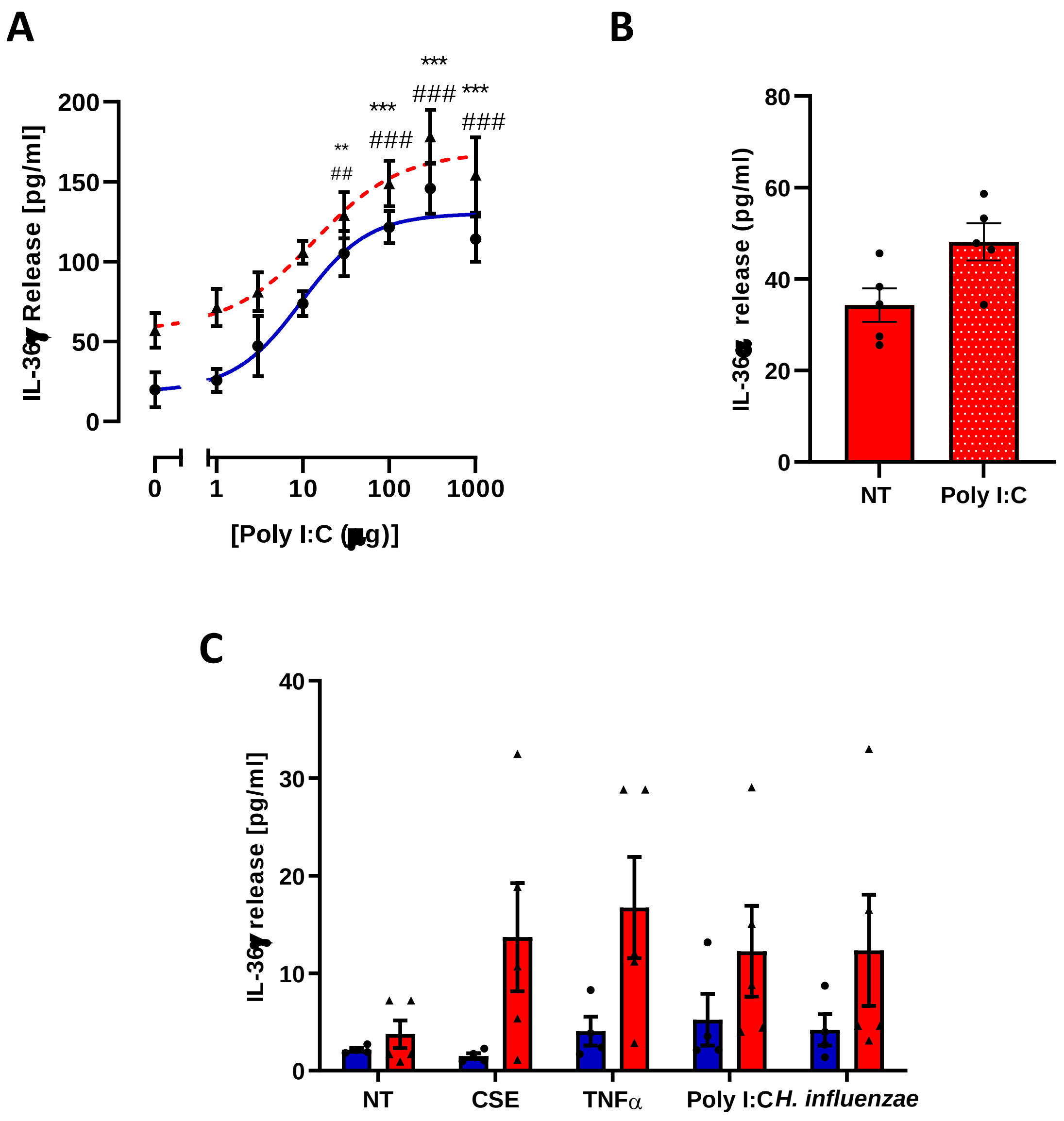


**Supplementary Figure 3. Poly I:C induces IL-36γ release from SAEC, but not SAF, whilst not inducing IL-36α.** A) Small airway epithelial cells from non-smokers (NS, ●, blue n=3) and COPD patients (▲, red n=3) were exposed to increasing concentrations of poly I:C for 24h. Media was collected and IL-36γ was measured by ELISA. B) SAEC were treated with 100 μg/ml of Poly I:C for 24 hours and IL-36α release detected by ELISA. C) SAF from non-smokers (NS, ●, blue, n=4) and COPD patients (▲, red, n=5) were exposed to media alone (no treatment: NT), 10% (v/v) cigarette smoke extract (CSE), 10 ng/ml TNFα, 100μg/ml poly I:C, or 1.5x$10^{10}$ CFU/ml *H. influenzae* for 24h. Media was collected and IL-36γ release measured by ELISA.

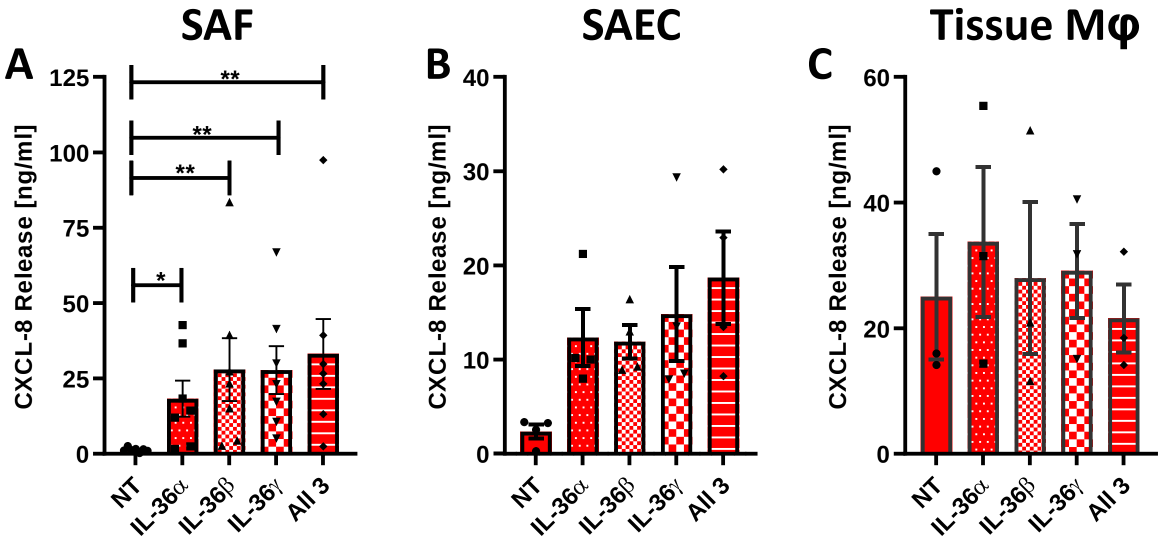


**Supplementary Figure 4. Effect of IL-36 cytokines alone and in combination on CXCL-8 release from small airway fibroblasts, human small airway epithelial cells and lung tissue macrophages.** A) Small airway fibroblasts, B) small airway epithelial cells and C) lung tissue macrophages were incubated in the absence (non-treated: NT) or presence of 33 ng/ml IL-36α, IL-36β, IL-36γ or all three in combination for 24h. Media was harvested, and release of CXCL-8 measured by ELISA. Data are means ± SEM and analysed by Kruskal-Wallis test with post-hoc Dunn's test; * P <0.05, ** P<0.01.

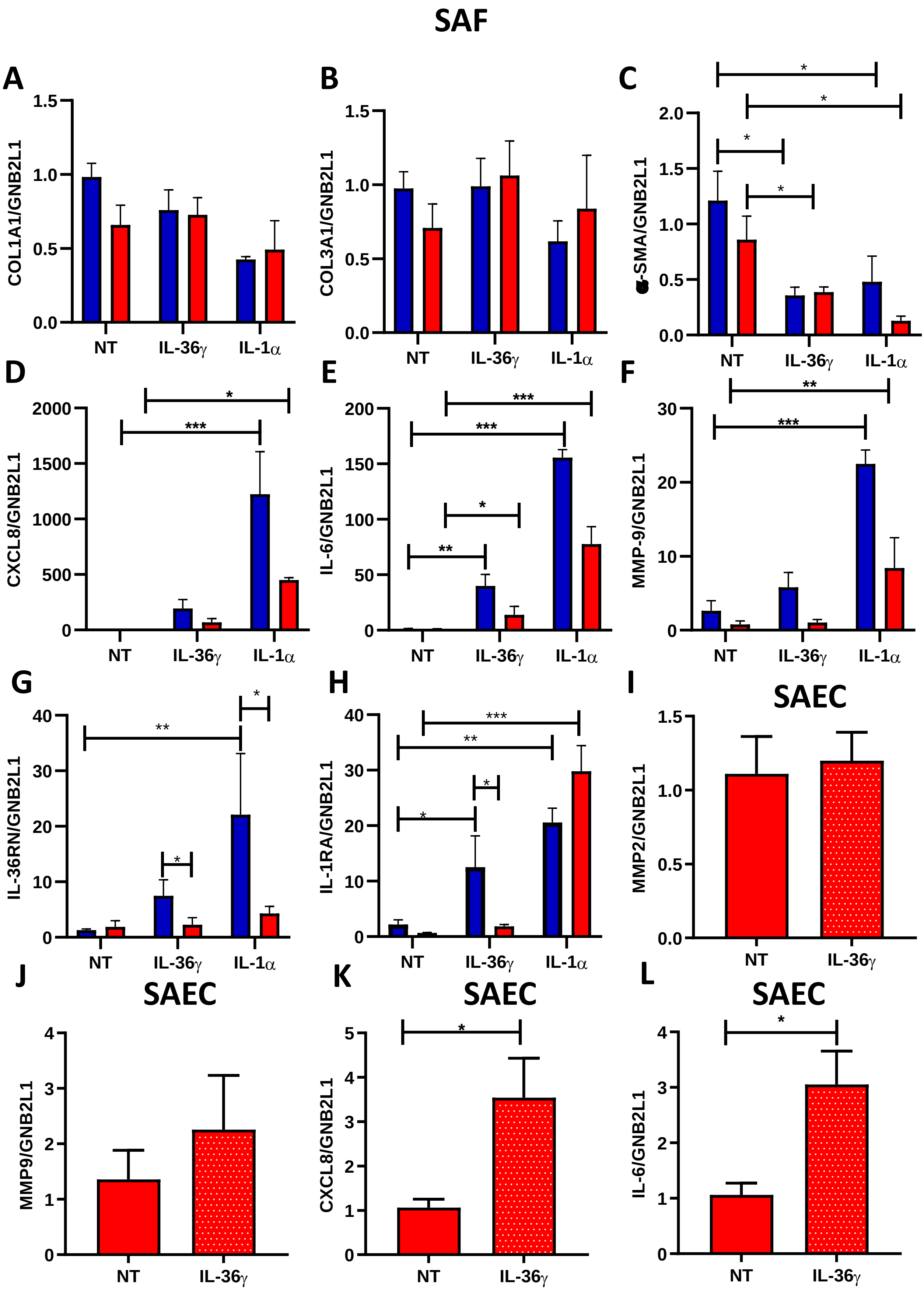
SAF
A
COL1A1/GNB2L1
1.5
1.0
0.5
0.0
NT
IL-36γ
IL-1α
B
COL3A1/GNB2L1
1.5
1.0
0.5
0.0
NT
IL-36γ
IL-1α
C
α-SMA/GNB2L1
2.0
1.5
1.0
0.5
0.0
*
*
*
*
NT
IL-36γ
IL-1α
D
CXCL8/GNB2L1
2000
1500
1000
500
0
*
***
NT
IL-36γ
IL-1α
E
IL-6/GNB2L1
200
150
100
50
0
***
***
*
**
NT
IL-36γ
IL-1α
F
MMP-9/GNB2L1
30
20
10
0
**
***
NT
IL-36γ
IL-1α
G
IL-36RN/GNB2L1
40
30
20
10
0
*
**
*
NT
IL-36γ
IL-1α
H
IL-1RA/GNB2L1
40
30
20
10
0
***
**
*
*
NT
IL-36γ
IL-1α
I
SAEC
MMP2/GNB2L1
1.5
1.0
0.5
0.0
NT
IL-36γ
J
SAEC
MMP9/GNB2L1
4
3
2
1
0
NT
IL-36γ
K
SAEC
CXCL8/GNB2L1
5
4
3
2
1
0
*
NT
IL-36γ
L
SAEC
IL-6/GNB2L1
4
3
2
1
0
*
NT
IL-36γ

**Supplementary Figure 5. Effect of IL-36γ and IL-1α cytokines on gene expression in small airway fibroblasts (SAF) and small airway epithelial cells (SAEC).** Small airway fibroblasts from non-smokers (n=7, blue) or COPD (n=7, red) were treated with 100 ng/ml of IL-36γ or IL-1α ng/ml for 24h. RNA was collected and gene expression of A) COL1A1, B) COL3A1, C) α-SMA and D) CXCL8, E) IL-6, F) MMP-9, G) IL-36RN and H) IL-1RA were detected by qRT-PCR. Small airway epithelial cells (n=4) were treated with 100 ng/ml of IL-36γ for 24h. RNA collected and gene expression of I) MMP-2, J) MMP-9, K) CXCL8, and L) IL-6, were detected by qRT-PCR Data are means ± SEM and analysed by two way anova with post-hoc Dunnett's multiple comparisons test; * P <0.05, ** P<0.01, *** P<0.001.

**Supplementary Table 1. The characteristics of subjects for bronchoscopy**

| | **Non-smoker** | **Smoker** | **COPD** |
|---|---|---|---|
| Sex (F:M) | 2:8 | 3:7 | 5:5 |
| Age (years) | 62±10.6 | 56±6.2 | 64.5±13.2 |
| Smoking history (pack-years) | 0 | 27.8±10 | 37.8±14.6 |
| $FEV_1$ (L) | 3.2±0.5 | 2.9±0.7 | 1.6±0.5* |
| $FEV_1$ (% predicted normal) | 100.2±10.0 | 86.6±15.9 | 62.9±22.7** |
| FVC (L) | 3.9±0.75 | 3.9±1.0 | 2.9±0.75 |
| $FEV_1$/FVC | 0.79±0.1 | 0.75±0.1 | 0.52±0.1*** |

Abbreviations: COPD = chronic obstructive pulmonary disease; $FEV_1$ = forced expiratory volume in one second; FVC = forced vital capacity; Data are expressed as mean value ± standard deviation and by Kruskal-Wallis test with post-hoc Dunn's test; * P <0.05, **P<0.01 and *** P<0.001 compared to non-smokers.

**Supplementary Table 2. The characteristics of study subjects for sputum samples**

| | Non-smoker | Smoker | COPD |
|---|---|---|---|
| **Sex (F:M)** | 9:9 | 4:4 | 7:13 |
| **Age (years)** | 46.25±11.05 | 55.8±3.2 | 68.6±9.7** |
| **Smoking history (Pack-years)** | 0 | 37±8.6*** | 44.7±31.6*** |
| **$FEV_1$ (L)** | 2.9±0.9 | 2.67±0.5 | 1.46±0.7*** |
| **$FEV_1$ (% predicted normal)** | 97.4±17.3 | 95.8±13.3 | 55.6.1±26.5*** |
| **FVC (L)** | 3.7±1.2 | 2.6±0.5 | 3.1±0.9 |
| **$FEV_1$/FVC** | 0.78±0.7 | 0.77±0.1 | 0.46±0.2*** |

Abbreviations: COPD = chronic obstructive pulmonary disease; $FEV_1$ = forced expiratory volume in one second; FVC = forced vital capacity. Data are expressed as mean value ± standard deviation and by Kruskal-Wallis test with post-hoc Dunn's test; **P<0.01 and *** P<0.001 compared to non-smokers.

**Supplementary Table 3. The characteristics of study subjects for nasal absorption samples**

| | Non-smoker | COPD |
|---|---|---|
| **Sex ratio (F:M)** | 6:2 | 8:12 |
| **Age (years)** | 61.1±3.2 | 66.5±2.1 |
| **Smoking History (pack years)** | 0.0±0.0 | 51.47±7.5*** |
| **$FEV_1$ (L)** | 2.8±0.3 | 1.4±0.6** |
| **$FEV_1$ % Predicted** | 109.4±5.5 | 54.75±4.3*** |
| **FVC (L)** | 3.8±0.5 | 3.0±1.9 |
| **$FEV_1$:FVC** | 0.75±0.0 | 0.48±0.0*** |

Abbreviations: COPD = chronic obstructive pulmonary disease; $FEV_1$ = forced expiratory volume in one second; FVC = forced vital capacity. Data are expressed as mean value ± standard deviation and analyzed by Mann-Whitney U test; * P <0.05 ***P<0.001 compared to non-smokers.

**Supplementary Table 4. The characteristics of study subjects for primary tissue macrophages**

| | Non-smoker | Smoker | COPD |
|---|---|---|---|
| **Sex (F:M)** | 7:3 | 10:14 | 13:16 |
| **Age (years)** | 69±7.7 | 66±12.2 | 69±8.9 |
| **Smoking history (Pack-years)** | 0 | 35.1±27.8*** | 38.6±18.1** |
| **$FEV_1$ (L)** | 2.2±0.7 | 2.5±0.6 | 1.2±0.6* |
| **$FEV_1$ (% predicted normal)** | 92.2±24.4 | 103.2±22.5 | 52.1.1±29.6** |
| **FVC (L)** | 2.6±0.8 | 3.5±0.8 | 2.7±1.1 |
| **$FEV_1$/FVC** | 0.8±0.1 | 0.72±0.1 | 0.43±0.1*** |

Abbreviations: COPD = chronic obstructive pulmonary disease; $FEV_1$ = forced expiratory volume in one second; FVC = forced vital capacity. Data are expressed as mean value ± standard deviation and by Kruskal-Wallis test with post-hoc Dunn's test; * P <0.05, **P<0.01 and *** P<0.001 compared to non-smokers.

**Supplementary Table 5. The characteristics of study subjects for primary airway epithelial cells**

| | Non-smoker | COPD |
|---|---|---|
| **Sex ratio (F:M)** | 7:4 | 10:7 |
| **Age (years)** | 71.7±4.3 | 68.9±6.5 |
| **Smoking History (pack years)** | 0.0±0.0 | 41.3±13.2** |
| **$FEV_1$ (L)** | 2.1±0.6 | 1.1±0.89* |
| **$FEV_1$ % Predicted** | 95.5±19.3 | 50.6±25.8** |
| **FVC (L)** | 2.8±0.7 | 2.5±0.8 |
| **$FEV_1$:FVC** | 0.74±0.1 | 0.5±0.2** |

Abbreviations: COPD = chronic obstructive pulmonary disease; $FEV_1$ = forced expiratory volume in one second; FVC = forced vital capacity; Data are expressed as mean value ± standard deviation and analyzed by Mann-Whitney U test; * $P < 0.05$ ** $P < 0.01$ compared to non-smokers.

**Supplementary Table 6. The characteristics of study subjects for primary small airway fibroblasts**

| | Non-smoker | COPD |
|---|---|---|
| **Sex ratio (F:M)** | 7:9 | 7:5 |
| **Age (years)** | 64.6±9.1 | 65.5±8.7 |
| **Smoking History (pack years)** | 0.0±0.0 | 34.5±18.3** |
| **$FEV_1$ (L)** | 2.7±0.9 | 1.5±0.8* |
| **$FEV_1$ % Predicted** | 92.7±11.6 | 52.9±25.6*** |
| **FVC (L)** | 3.4±0.8 | 3.2±1.1 |
| **$FEV_1$:FVC** | 0.79±0.1 | 0.54±0.2** |

Abbreviations: COPD = chronic obstructive pulmonary disease; $FEV_1$ = forced expiratory volume in one second; FVC = forced vital capacity; Data are expressed as mean value ± standard deviation and analyzed by Mann-Whitney U test; * P <0.05, **P<0.01 and *** P<0.001 compared to non-smokers

**Supplementary Table 7. The characteristics of study subjects for lung homogenate samples**

| | **Non-smokers (n=9)** | **Smokers (n=9)** | **GOLD Stage 1 (n=9)** | **GOLD Stage 2 (n=9)** | **GOLD Stage 3 (n=3)** | **GOLD Stage 4 (n=6)** |
|---|---|---|---|---|---|---|
| **Age** (years) | 63.4±13.6 | 63±12.3 | 67.7±7.0 | 63.0±9.3 | 62.3±10.9 | 59.8±4.5 |
| **Sex** (F:M) | 7:2 | 5:4 | 3:5 | 5:4 | 1:2 | 4:2 |
| **$FEV_1$** (L) | 2.56±0.6 | 2.8 ±0.6 | 2.7±0.6 | 1.8±0.4* | 1.69±0.4* | 0.5±0.18** |
| **$FEV_1$** (% predicted) | 97.2±16.4 | 99.4±13.3 | 89.1±3.9 | 65.4±17.5** | 49.7±4.3* | 16.1±2.9*** |
| **FVC** (L) | 3.2±1.1 | 3.6±0.9 | 4.0±0.9 | 3.1±0.8 | 3.0.1±0.7 | 1.75±0.5* |
| **$FEV_1$:FVC** | 80.3±4.9 | 75.4±4.1 | 64.3±3.6* | 61.5±7.9** | 51.8±7.1* | 28.5±8.1*** |
| **Smoking History** (Pack years) | 0±0 | 61.1±32.4 | 44.3±17.0 | 57.7±35.4 | 46.6±21.8 | 38.6±15.9 |

Abbreviations: COPD = chronic obstructive pulmonary disease; $FEV_1$ = forced expiratory volume in one second; FVC = forced vital capacity. Data are expressed as mean value ± standard deviation and by Kruskal-Wallis test with post-hoc Dunn's test; * P <0.05, **P<0.01 and *** P<0.001 compared to non-smokers.
